# Neural hierarchical models of ecological populations

**DOI:** 10.1101/759944

**Authors:** Maxwell B. Joseph

**Affiliations:** Earth Lab, Cooperative Institute for Research in Environmental Sciences, University of Colorado Boulder, Boulder, CO 80303, USA

## Abstract

Neural networks are increasingly being used in science to infer hidden dynamics of natural systems from noisy observations, a task typically handled by hierarchical models in ecology. This paper describes a class of hierarchical models parameterized by neural networks: neural hierarchical models. The derivation of such models analogizes the relationship between regression and neural networks. A case study is developed for a neural dynamic occupancy model of North American bird populations, trained on millions of detection/non-detection time series for hundreds of species, providing insights into colonization and extinction at a continental scale. Flexible models are increasingly needed that scale to large data and represent ecological processes. Neural hierarchical models satisfy this need, providing a bridge between deep learning and ecological modeling that combines the function representation power of neural networks with the inferential capacity of hierarchical models.

## Introduction

Deep neural networks have proved useful in myriad tasks due their ability to represent complex functions over structured domains (LeCun *et al*. 2015). While ecologists are beginning to use such approaches, e.g., to identify plants and animals in images (Norouzzadeh *et al*. 2018; Fricker *et al*. 2019), there has been relatively little integration of deep neural networks with ecological models.

Ecological processes are difficult to observe. Inference often proceeds by modeling the relationship between imperfect data and latent quantities or processes of interest with hierarchical models (Wikle 2003). For example, occupancy models estimate the presence or absence of a species using imperfect detection data (MacKenzie *et al*. 2002), and “dynamic” occupancy models estimate extinction and colonization dynamics (MacKenzie *et al*. 2003). Population growth provides another example, motivating hierarchical models that link noisy observations to mechanistic models (De Valpine & Hastings 2002). In such models, it is often desirable to account for heterogeneity among sample units (e.g., differences among habitats and survey conditions), to better understand ecological dynamics.

Many hierarchical models in ecology account for heterogeneity among sample units using a linear combination of explanatory variables, despite there often being reasons to expect non-linearity (Lek *et al*. 1996; Austin 2002; Oksanen & Minchin 2002). A variety of solutions exist to account for non-linearity. For example, Gaussian processes have been used for species distribution models (Latimer *et al*. 2009; Golding & Purse 2016), in animal movement models (Johnson *et al*. 2008), and in point process models for distance sampling data (Johnson *et al*. 2010; Yuan *et al*. 2017). Generalized additive models also have been used to account for spatial autocorrelation (Miller *et al*. 2013; Webb *et al*. 2014), nonlinear responses to habitat characteristics (Knapp *et al*. 2003; Bled *et al*. 2013), and differential catchability in capture-recapture studies (Zwane & Van der Heijden 2004).

Machine learning provides additional tools for approximating nonlinear functions in hierarchical models. For example, Hutchinson *et al*. (2011) combined a site-occupancy model with an ensemble of decision trees to predict bird occurrence, combining a structured observation and process model from ecology with a flexible random forest model. Similar hybrid approaches could be developed for other classes of ecological models and/or machine learning methods. Neural networks seem particularly worthy of attention, given their success in other domains (LeCun *et al*. 2015).

Neural networks have been used in ecology, but to date have not been integrated into hierarchical models that explicitly distinguish between ecological processes and imperfect observations. For example, neural networks have modeled observed abundances of aquatic organisms, but without distinguishing observed from true abundance (Chon *et al*. 2001; Jeong *et al*. 2001, 2008; Malek *et al*. 2012). Neural networks also have been used to model stock-recruitment and apparent presence/absence data (Manel *et al*. 1999; Chen & Hare 2006; Özesmi *et al*. 2006; Harris 2015; Chen *et al*. 2016), but have not yet been extended to account for imperfect detection, despite increasing recognition of its importance (Guillera-Arroita 2017; Tobler *et al*. 2019). Notably, standard neural networks that classify or predict data are of limited use for ecological applications where imperfect data are used to learn about dynamics of hard to observe systems. As a solution, this paper describes neural hierarchical models that combine the function representation capacity of neural networks with hierarchical models that represent ecological processes.

### Related work

#### Neural networks

Neural networks are function approximators. Linear regression is a special case, where an input vector **x** is mapped to a predicted value *y*:

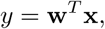

where **w** is a parameter vector (Fig. 1a). Predicted values are linear combinations of the inputs **x** due to the product **w**^*T*^ **x**, which restricts the complexity of the function mapping **x** to *y*.

**Figure 1:**
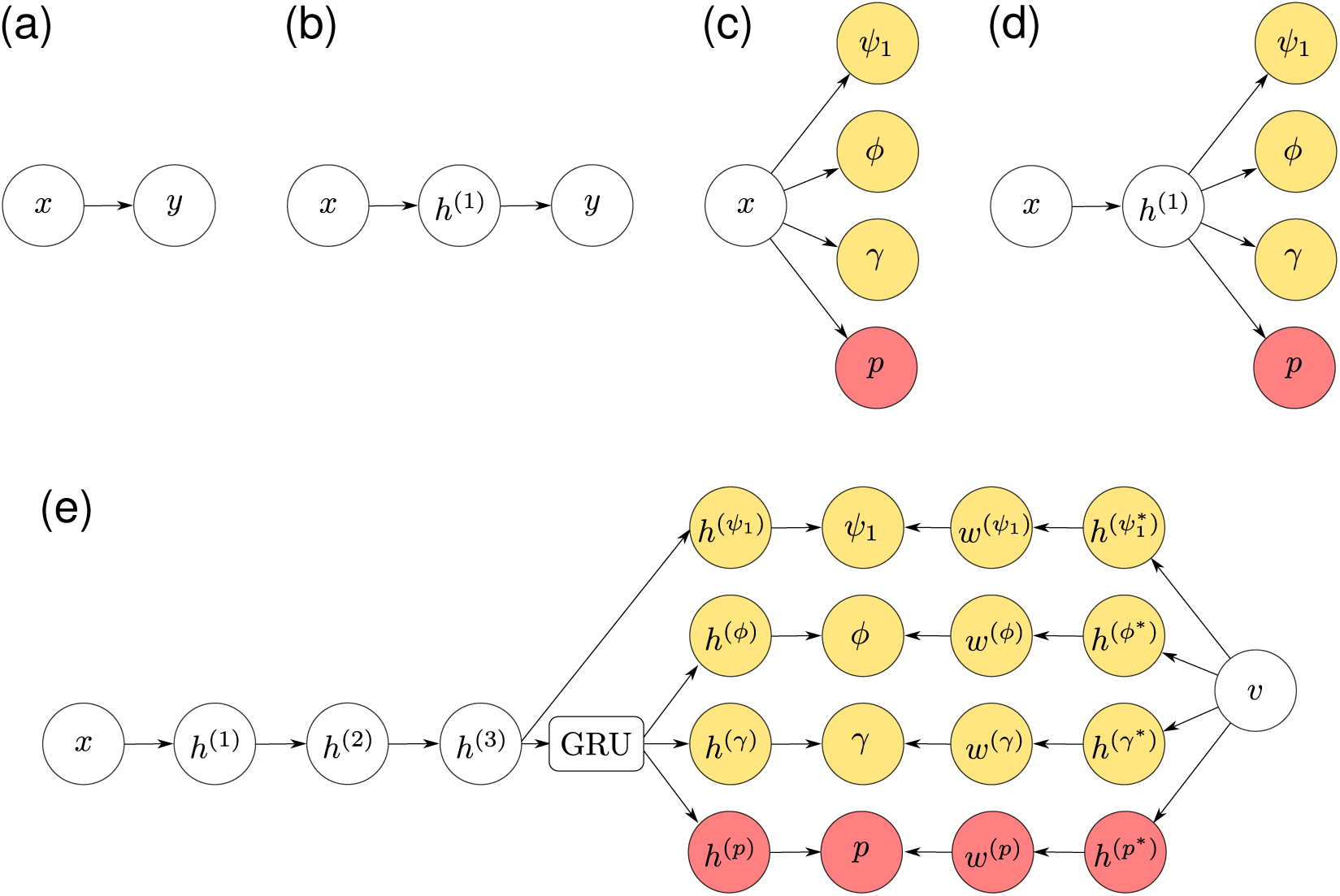
Computation graphs for (a) linear regression, (b) a neural network with one hidden layer, (c) a dynamic occupancy model, (d) a single species neural dynamic occupancy model, and (e) a deep multi-species neural dynamic occupancy model. Yellow and red indicate quantities specific to process and observation components respectively. Inputs are represented by *x*, and predicted values in panels (a) and (b) by *y*. Hidden layers are represented by *h*, with layer-specific superscripts. Outputs include initial occupancy (*ψ*_1_), persistence (*ϕ*), colonization (*γ*), and detection (*p*) probabilities. Latent species embedding vectors are represented by *v*, and GRU indicates a gated recurrent unit.

Instead of modeling outputs as linear functions of **x**, a model can be developed that is a linear function of a nonlinear transformation of **x**. Nonlinear transformations can be specified via polynomial terms, splines, or another basis expansion (Hefley *et al*. 2017), but neural networks parameterize the transformation via a set of sequential “hidden layers” (Goodfellow *et al*. 2016).

In a neural network with one hidden layer, the first hidden layer maps the length *D* input **x** to a length *D*^(1)^ vector of “activations” **a**^(1)^ = **W**^(1)^**x**, where **W**^(1)^ is a *D*^(1)^ × *D* parameter matrix. The activations are passed to a differentiable nonlinear activation function *g* to obtain the “hidden units” of the first layer **h** ^(1)^ = *g*(**a**^(1)^) (Fig. 1b).

The final layer of a neural network maps the a hidden layer to an output. For a neural network with one hidden layer, if the output variable **y** is a *K* dimensional vector, the output unit activations are given by **a**^(2)^ = **W**^(2)^**h**^(1)^, where **W**^(2)^ is a *K* × *D*^(1)^ parameter matrix. The output activation can be written as a composition of functions that transform the inputs *x*:

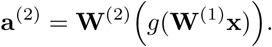

Similar to link functions in generalized linear models, outputs can be transformed by an output activation function. For example, if **y** is unbounded, the identity function can act as an output activation so that **y** = **a**^(2)^. Neural networks that predict probabilities typically use a sigmoid (inverse logit) activation function.

Neural networks usually are trained using stochastic gradient based optimization to minimize a loss function, e.g., the negative log likelihood of a Gaussian distribution for a regression task, a Bernoulli distribution for binary classification, or a Poisson distribution for a count model. Partial derivatives of the model parameters are computed with respect to the loss via backpropagation, and parameters are updated to reduce the loss. In practice, these partial derivatives are often computed via automatic differentiation over a “mini-batch” of samples, which provides a noisy estimate of the gradient (Ruder 2016).

Neural networks are popular both because of their practical successes in a wide variety of applications, and because they possess some desirable theoretical properties. A neural network with suitable activation functions and a single hidden layer containing a finite number of neurons can approximate nearly any continuous function on a compact domain (Cybenko 1989; Hornik 1991). “Deep” neural networks with many sequential hidden layers also act as function approximators (Lu *et al*. 2017). The field of deep learning, which applies such networks, provides a variety of network architectures to account for temporal structure (Hochreiter & Schmidhuber 1997), spatial structure on regular grids (Long *et al*. 2015), or graphs (Niepert *et al*. 2016), sets of unordered irregular points (Li *et al*. 2018), and spatiotemporal data on grids or graphs (Xingjian *et al*. 2015; Jain *et al*. 2016).

The potential for deep learning has been recognized in Earth science (Reichstein *et al*. 2019), the natural sciences (Ching *et al*. 2018; Gazestani & Lewis 2019; Roscher *et al*. 2019), physical sciences (Carleo *et al*. 2019), chemical sciences (Butler *et al*. 2018), and ecology (Christin *et al*. 2018; Desjardins-Proulx *et al*. 2019). For example, models of lake temperature that combine neural networks with loss functions consistent with known physical mechanisms perform better than physical models and neural networks applied alone (Karpatne *et al*. 2017). Similarly, generative adversarial networks with loss functions that encourage mass-balance have expedited electromagnetic calorimeter data generation from the Large Hadron Collider (Paganini *et al*. 2018; Radovic *et al*. 2018). Convolutional neural networks also have been successfully deployed in population genetics to make inferences about introgression, recombination, selection, and population sizes (Flagel *et al*. 2018). There are various ways to combine science knowledge with deep learning. Ba *et al*. (2019) provide a useful taxonomy in the context of physics-based deep learning. In ecology, hierarchical models present an opportunity to build upon existing approaches to derive science-based deep learning methods.

#### Hierarchical models

Hierarchical models combine a data model, a process model, and a parameter model (Berliner 1996; Wikle 2003). Data models represent the probability distribution of observations conditioned on a process and some parameters, e.g., the probability of capturing a marked animal, given its true state (alive or dead). Process models represent states and their dynamics, conditioned on some parameters. State variables are often incompletely observed, e.g., whether an individual animal is alive or whether a site is occupied. Parameter models represent probability distributions for unknown parameters - priors in a Bayesian framework. In a non-Bayesian setting, parameters are treated as unknowns and estimated from the data, but parameter uncertainty is not represented using probability distributions (Cressie *et al*. 2009).

### Neural hierarchical models

Neural hierarchical models are hierarchical models in which the observation, process, or parameter model is parameterized by a neural network. These models are hierarchical (sensu Berliner (1996)) if they distinguish between a modeled process and available data, e.g., between partial differential equations and noisy observations of their solutions (Raissi 2018). “Deep Markov models” - hidden Markov models parameterized by neural networks - provide an example, with successful applications in polyphonic music structure discovery, patient state reconstruction in medical data, and time series forecasting (Krishnan *et al*. 2017; Rangapuram *et al*. 2018). State-space neural networks that use recurrent architectures provide another example dating back two decades (Zamarreño & Vega 1998; Van Lint *et al*. 2002). This class of models inherits the flexibility and scalability of neural networks, along with the inferential power of hierarchical models, but applications in ecology and environmental science are only just beginning to emerge (Wikle 2019).

Construction of such models from existing hierarchical models is straightforward. For example, one can propose neural variants of occupancy models (MacKenzie *et al*. 2002), dynamic occupancy models (MacKenzie *et al*. 2003; Royle & Kéry 2007), N-mixture models (Royle 2004), mark-recapture models (Jolly 1965; Calvert *et al*. 2009), and other hidden Markov models (Patterson *et al*. 2009, 2017; Langrock *et al*. 2012). Output activation functions can be determined from inverse link functions, e.g., sigmoid (inverse logit) activations for probabilities, and loss functions can be constructed from the negative log likelihoods (see Appendix S1 in Supporting Information for example model specifications). Specialized neural network architectures that operate on structured data can be readily integrated into such models. For example, Appendix S2 in Supporting Information provides a simulated animal movement case study where aerial imagery is mapped to state transition probabilities of a hidden Markov model with a convolutional neural network. To provide an empirical use case, a neural dynamic occupancy model is developed for extinction and colonization dynamics of North American bird communities.

#### Case study

The North American Breeding Bird Survey (BBS) is a large-scale annual survey aimed at characterizing trends in roadside bird populations (Link & Sauer 1998; Sauer & Link 2011; Sauer *et al*. 2013; Pardieck *et al*. 2018). Thousands of routes are surveyed once a year during the breeding season. Surveys consist of volunteer observers that stop 50 times at points 800 meters apart on a transect, recording all birds detected within 400 meters for three minutes. Species can be present, but not detected, and may go locally extinct or colonize new routes from year to year, motivating the development of dynamic occupancy models which use imperfect detection data to estimate latent presence or absence states (MacKenzie *et al*. 2003; Royle & Kéry 2007).

This case study uses BBS data from 1997-2018 excluding unidentified or hybrid species, restricting the analysis to surveys meeting the official BBS criteria (Pardieck *et al*. 2018). The resulting data consists of 647 species sampled at 4,540 routes, for a total of 59,384 surveys (not every route is surveyed in each year), 2,937,380 observation history time series, and 38,421,448 detection/non-detection observations.

#### Process model

A multi-species dynamic occupancy model for spatially referenced routes *s* = 1, …, *S*, surveyed in years *t* = 1, …*T*, for species *j* = 1, …, *J* aims to estimate colonization and extinction dynamics through time. The true occupancy state *z*_*t,s,j*_ = 1 if species *j* is present, and *z*_*t,s,j*_ = 0 if species *j* is absent. The model represents *z*_*t,s,j*_ as a Bernoulli distributed random variable, where Pr(*z*_*t,s,j*_ = 1) = *ψ*_*t,s,j*_. The probability of occurrence on the first timestep is *ψ*_*t*=1,*s,j*_. Subsequent dynamics are determined by probabilities of persistence from time *t* to *t* + 1 denoted *ϕ*_*t,s,j*_, and probabilities of colonization from time *t* to *t* + 1 denoted *γ*_*t,s,j*_, so that the probability of occurrence in timesteps *t* = 2, …, *T* is (MacKenzie *et al*. 2003):

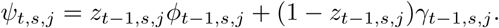

#### Observation model

Let *y*_*t,s,j*_ represent the number of stops where species *j* was detected in year *t* on route *s*. Conditional on a species being present, it is detected at each stop with probability *p*_*t,s,j*_. Assume that there are no false-positive detections, so that if a species is absent, it cannot be detected (but see Royle & Link 2006). With *k* = 50 replicate stops on each transect, the observations can be modeled using a Binomial likelihood for one species-route-year combination:

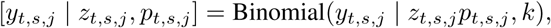

where square brackets denote the probability function (e.g., in this case, [*y*_*t,s,j*_ | *z*_*t,s,j*_, *p*_*t,s,j*_] = Pr(*Y*_*t,s,j*_ = *y*_*t,s,j*_ | *z*_*t,s,j*_, *p*_*t,s,j*_) where *Y*_*t,s,j*_ is a discrete random variable and *y*_*t,s,j*_ is a particular value). The joint likelihood corresponds to the product of these terms for all years, routes, and species (Dorazio *et al*. 2010).

#### Parameter models

Heterogeneity in parameter values among routes, years, species, and surveys was modeled using three different approaches.

1. **Single-species baseline models** mapped input features to occupancy parameters with a linear combination on the logit scale (MacKenzie *et al*. 2003) (Fig. 1c).
2. **Single-species neural hierarchical models** mapped inputs to occupancy parameters using a neural network with one hidden layer (Fig. 1d).
3. **A multi-species deep neural hierarchical model** was developed to model occupancy dynamics of all species simultaneously (Fig. 1e).

Input features included Environmental Protection Agency (EPA) level one ecoregions, the first eight principal components of the standard 19 WorldClim Bioclimatic variables averaged across the years 1970-2000 (Fick & Hijmans 2017), BBS route spatial coordinates, distance from the coast, elevation, and road density within a 10 km buffer of each route (Meijer *et al*. 2018). The model of detection probabilities additionally included survey-level features including temperature, duration, wind, and air conditions.

The neural hierarchical models were motivated by joint species distribution models in which species load onto a shared set of latent factors (Thorson *et al*. 2015, 2016; Warton *et al*. 2015; Ovaskainen *et al*. 2016; Tikhonov *et al*. 2017), and by recent work on deep neural basis expansions (McDermott & Wikle 2019; Wikle 2019). Analogously, the neural networks combined inputs into a latent vector for each route (the hidden layers), which act as latent factors that are mapped to parameters (Fig. 1d). Because the single species neural hierarchical models were fit separately for each species, latent factors were species-specific. In contrast, latent factors were shared among species in the multi-species model.

The multi-species neural dynamic occupancy model additionally built upon previous work on deep multi-species embedding. Deep multi-species embedding uses vector-valued “entity embeddings” to represent each species (Chen *et al*. 2016; Guo & Berkhahn 2016). These entity embeddings are mappings from categorical data (e.g., a species identity) to continuous numeric vector representations. Embeddings are used extensively in language models such as word2vec, which maps words to vector spaces (Mikolov *et al*. 2013).

Further, the multi-species model combined species embeddings with encoder-decoder components to estimate occupancy parameters. Encoder-decoder neural networks are used in sequence-to-sequence translation (Sutskever *et al*. 2014). They encode inputs (e.g., a sequence of words) into a latent vector space, then decode that vector representation to generate another sequence using a neural network. Similarly, the multi-species model encoded route-level features into vector representations. These route vectors were decoded by a recurrent neural network to generate a multivariate time series of latent vectors associated with colonization, persistence, and detection (Chung *et al*. 2014). Finally, the multi-species model combined these route-level latent vectors with species-level embeddings to compute colonization, persistence, and detection probabilities (Fig. 1e). For additional details, see Appendix S3 in Supporting Information.

#### Model comparisons

To compare the performance of the three modeling approaches, the data were partitioned into a training, validation, and test set at the EPA level two ecoregion level (Roberts *et al*. 2017). All routes within an ecoregion were assigned to the same partition. This resulted in 2154 training routes, 948 validation routes, and 1438 test routes. K-fold cross validation would also be possible, though it requires retraining of each model K times (Roberts *et al*. 2017). Because of the size of the BBS data and the computational resources required to train these models, a simpler train/validation/test split was used.

For each of the three modeling approaches, routes in the training set were used for parameter estimation. Single-species models were fit separately for each species (modeling approaches 1 and 2), and one multi-species model (approach 3) was fit using all of the training data. Then, using the trained models, the mean predictive log-density of the validation data was evaluated to identify the best performing model (Gelman *et al*. 2014). This step indicated which model fit best to the withheld validation data. Finally, the best performing model was retrained using the training and validation data, and its predictive performance was evaluated on the withheld test set (Russell & Norvig 2016).

#### Final model evaluation

The final model’s performance was evaluated quantitatively and qualitatively. Quantitative predictive performance was evaluated in two ways. First, 95% prediction interval coverage was computed for test set counts. Prediction intervals were constructed using the quantiles of the binomial distribution, marginalizing over latent occupancy states. Second, the area under the receiver operator characteristic curve (AUC) was computed for binarized test set counts. The AUC analysis also marginalized over latent states to derive predicted probabilities, e.g., Pr(*y*_*t,s,j*_ = 0 | *p*_*t,s,j*_, *ψ*_*t,s,j*_) = 1 − *ψ*_*t,s,j*_ + *ψ*_*t,s,j*_ (1 − *p*_*t,s,j*_)^50^, where the parameters *p* and *ψ* are estimated by the model. These two approaches provide information on how well the final model could predict the number of stops with detections at each route, and whether any detections would occur on each route, respectively.

Qualitative analyses were based on predicted occupancy states from the final model. The most likely occupancy states were computed at all BBS routes for each year and species using the Viterbi algorithm (Viterbi 1967). The estimated occupancy states were then used to compute finite sample population growth rates for each species (Royle & Kéry 2007). Occupancy state estimates were also used to compute annual spatial centroids for each species, by taking the spatial centroids of occupied route coordinates. These are hereafter referred to as “BBS range centroids” to differentiate from the actual centroid of a species’ entire range.

To evaluate whether the results were qualitatively consistent with previous findings about colonization and extinction gradients over species ranges, correspondence between BBS range centroids and both colonization and persistence probabilities were assessed using linear regression (Mehlman 1997; Doherty Jr *et al*. 2003; Royle & Kéry 2007). For this analysis, the response variable was colonization or persistence averaged over time, the predictor was distance from BBS range centroid. Survey routes were the sample units, and separate analyses were conducted for each species. To avoid bias associated with recently added BBS routes and variance associated with rare species, these analyses only used data from BBS routes surveyed in every year, species observed in every year, and species that occurred in 100 or more routes (Sauer *et al*. 2017).

Finally, visualizations were developed to graphically interpret the final model. First, route-level features were visualized using t-distributed stochastic neighbor embedding, which maps high dimensional vectors to lower dimensional representations (Maaten & Hinton 2008). In this lower dimensional space, routes with similar embeddings are close together, and routes with dissimilar embeddings appear distant (Rauber *et al*. 2016). Second, species’ loading vectors were compared in terms of cosine similarity, which measures the orientation of loading vectors in latent space. Species’ occupancy should be positively related if loading vectors are oriented similarly. This expectation was checked graphically by comparing estimated occupancy time series of species that had the most similar and most different loading vectors.

## Results

The multi-species neural hierarchical model performed best on the withheld validation routes. The difference in mean validation set negative log likelihood was 5.015 relative to the baseline model and 2.108 relative to the single-species neural hierarchical model. For the final model, 95% prediction interval coverage for observed counts at withheld test set routes was 93.6%, with a standard deviation of 1.5%, a minimum of 86.5%, and a maximum of 97.6%. The model also predicted whether species would be detected on test set routes fairly well, with a mean test set AUC of 0.953, an among-route standard deviation of 0.032, a minimum of 0.667, and a maximum of 0.993 (Fig. 2).

**Figure 2:**
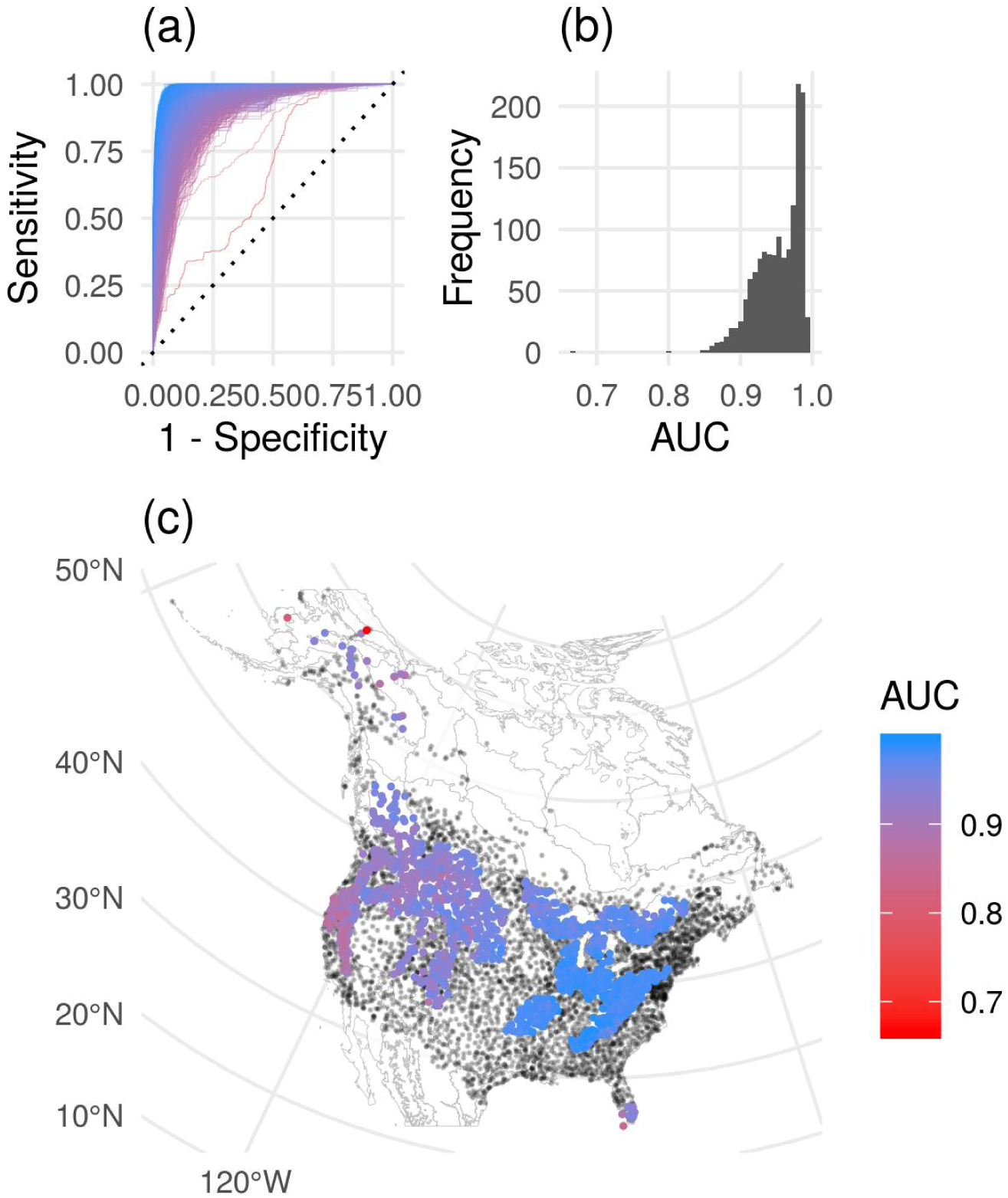
(a) Receiver operator characteristic curves of binarized detection/non-detection data for each survey route in the test set, colored by the area under the curve (AUC). Here, the x-axis is the complement of specificity: the ratio of the number of false positives (incorrectly predicted detections, with no observed detections) to the sum of false positives and true negatives (correctly predicted non-detections with observed non-detections). The y-axis is true positive rate: the ratio of the number of true positives (correctly predicted detections with observed detections) to the sum of true positives and false negatives (incorrectly predicted non-detections with observed detections). The overall distribution of AUC values is shown in (b), and (c) shows the locations of test set routes colored by AUC values, where black dots represent routes that were used to train the final model.

Qualitative results related to range shifts and population growth rates were plausible given previous work. The model identified the invasive Eurasian Collared-Dove (*Streptopelia decaocto*) as having the greatest range centroid displacement from 1997 to 2018 and the highest finite sample population growth rate (Fig. 3), consistent with its invasion of North America from Florida following its introduction in the 1980’s (Bled *et al*. 2011). Increasing trends also have previously been reported for the species with the next three highest population growth rates: Bald Eagle (*Haliaeetus leucocephalus*), Wild Turkey (*Meleagris gallopavo*), and Osprey (*Pandion haliaetus*) (Sauer *et al*. 2013).

**Figure 3:**
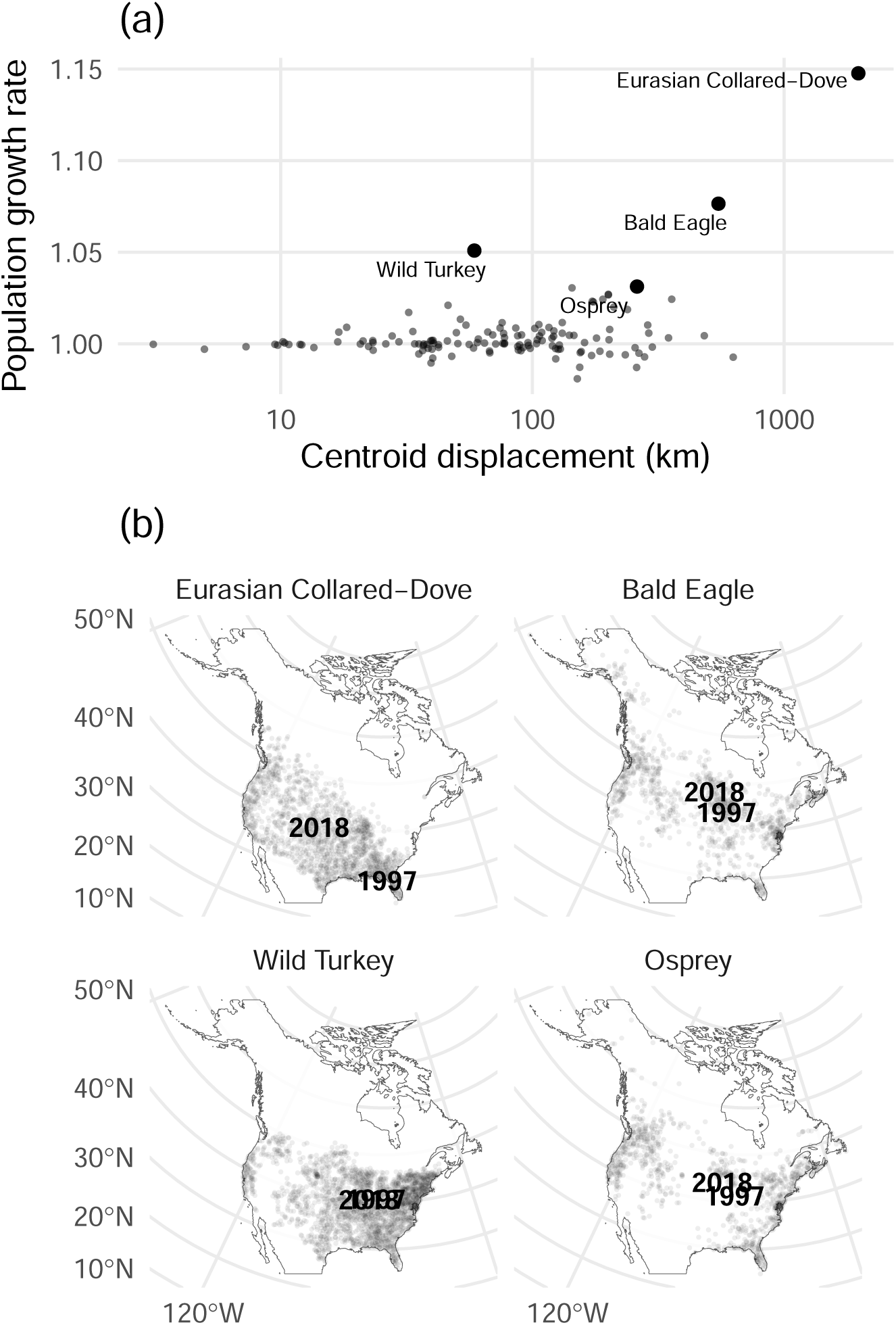
(a) Centroid displacement of bird species (x-axis) from 1997 to 2018 in kilometers vs. finite-sample population growth rate (y-axis), where a growth rate of 1 is stable, values less than one are decreasing, and values greater than one are increasing. Species with the highest population growth rates are highlighted. Panel (b) shows the locations of breeding bird survey range centroids in 1997 and 2018 for each highlighted species, along with grey points that represent survey routes where the species was estimated to be present in at least one year.

The majority (77%) of common species were less likely to persist at routes that were distant from their estimated BBS range centroids. Similarly, 98% of common species were less likely to colonize routes distant from their BBS range centroids. There were examples of species with positive and negative distance coefficients for persistence and colonization (Fig. 4a). Negative relationships were most apparent for common species that occupied a large fraction of BBS routes (Fig. 4b). Results for representative species are displayed in Fig. 4c-d.

**Figure 4:**
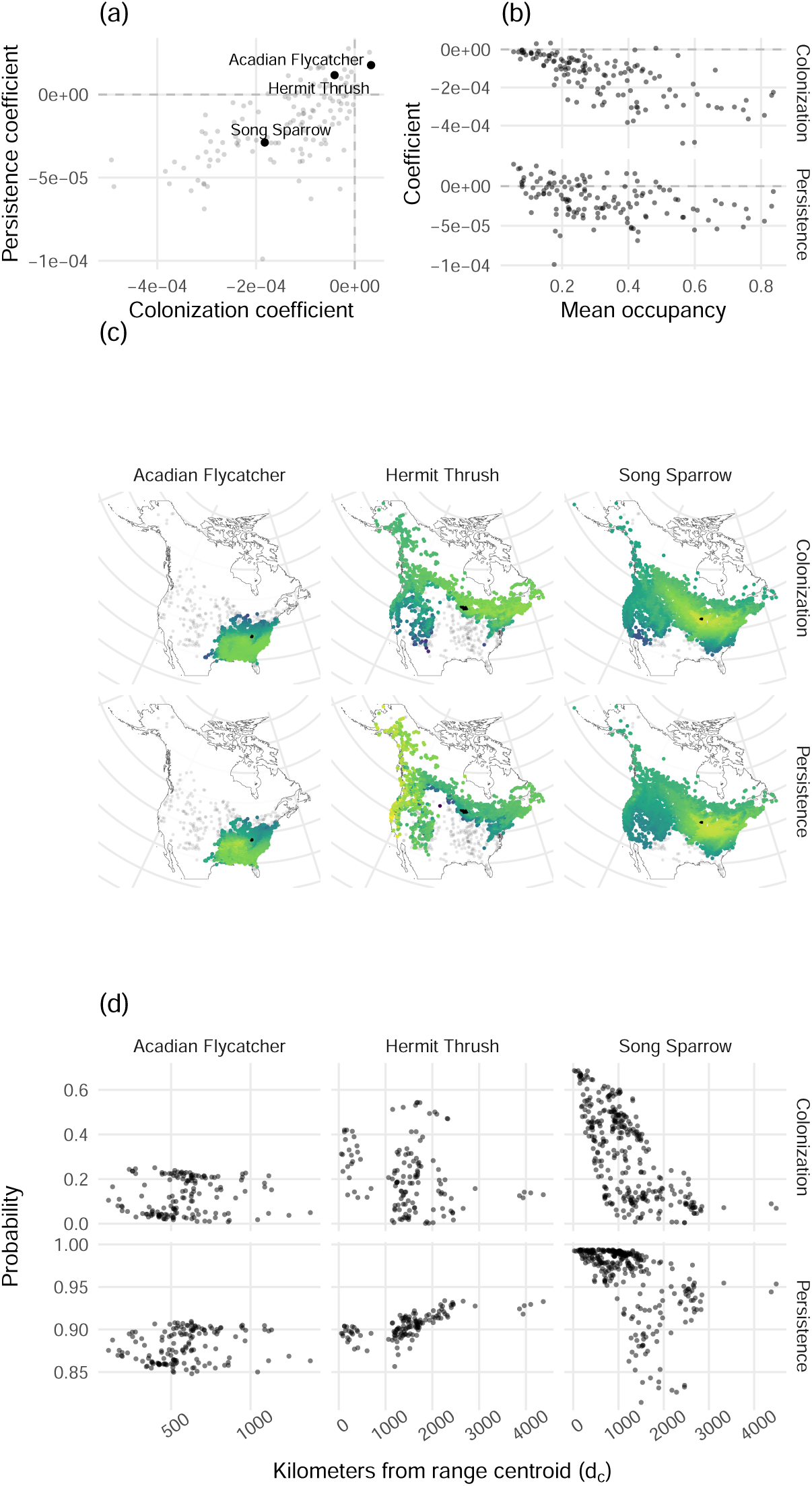
(a) Scatter plot relating species-specific distance decay coefficients for colonization and persistence, with focal species from each quadrant highlighted. (b) Mean finite-sample occupancy (x-axis) vs. coefficients relating distance from range centroid and the probability of colonization (y-axis, top row), and persistence (y-axis, bottom row). (c) Maps of colonization and persistence probabilities at each route where focal species were likely absent (in grey) and present (in color, normalized to increase the visibility of gradients). Centroids for each year are shown in solid black. (d) Colonization and persistence probabilities (y-axis) as a function of distance from range centroid, averaged among years.

Route vectors combined information from the categorical and continuous route-level features. Unsurprisingly (because ecoregion was an input feature), routes in the same ecoregions clustered together (Fig. 5a). Route embeddings also revealed relationships among ecoregions. For example, Marine West Coast Forest ecoregion routes were similar to Northwestern Forested Mountains routes, and most different from Northern Forests routes (Fig. 5a). Variation within clusters also related to continuous route-level features (Fig. 5b).

**Figure 5:**
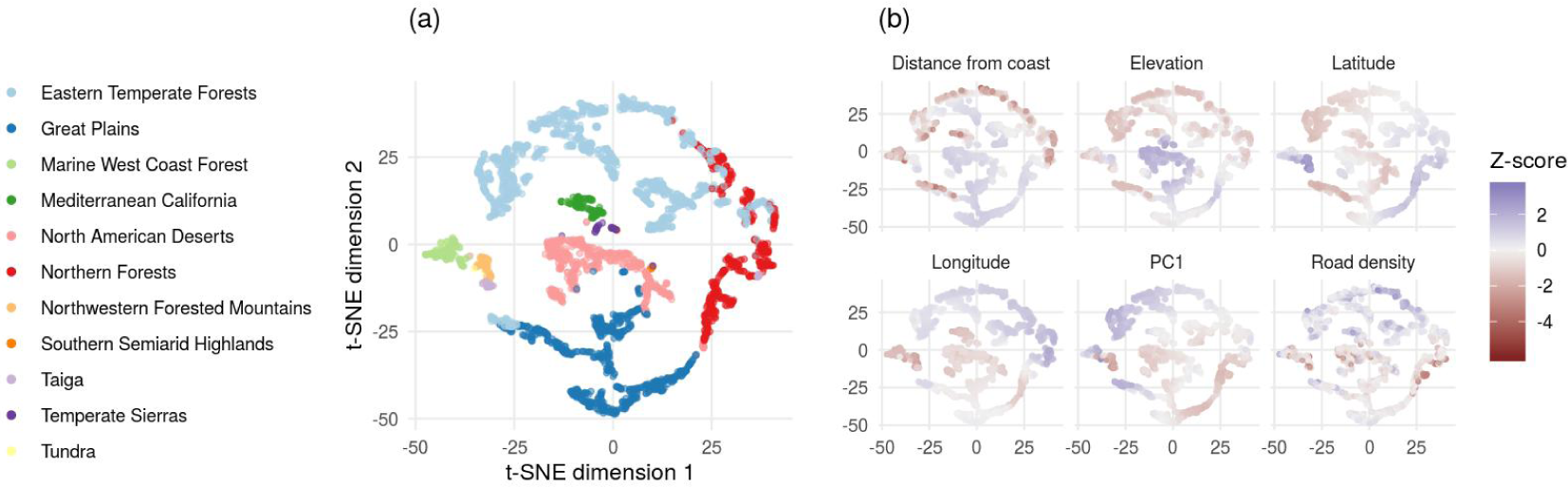
Clustering of North American Breeding Bird Survey routes by ecoregion and route-level features. All panels show a two dimensional plot of route vectors, computed using t-distributed stochastic neighbor embedding (t-SNE) on the latent vectors for initial occupancy, persistence, colonization, and detection probabilities. Each route is shown as a point. Color indicates (a) EPA level 1 ecoregion and (b) z-standardized continuous route-level features. PC1 refers to the first principal component of WorldClim climate data.

The model predicted non-linear dependence among species that is interpretable in terms of the cosine similarity among species-specific parameter vectors. For example, parameters for Mourning Dove (*Zenaida macroura*) were closest to those of Barn Swallow (*Hirundo rustica*), and most different from (Myrtle Warbler) Yellow-rumped Warbler (*Setophaga coronata coronata*) (Fig. 6a). On BBS routes where Mourning Doves are likely to occur, Barn Swallow are also likely to occur, and (Myrtle Warbler) Yellow-rumped Warbler are unlikely to occur. Species pairs of the most similar and most dissimilar loadings are also provided for Eurasian Collared-Dove and Bald Eagle (Fig. 6b-c).

**Figure 6:**
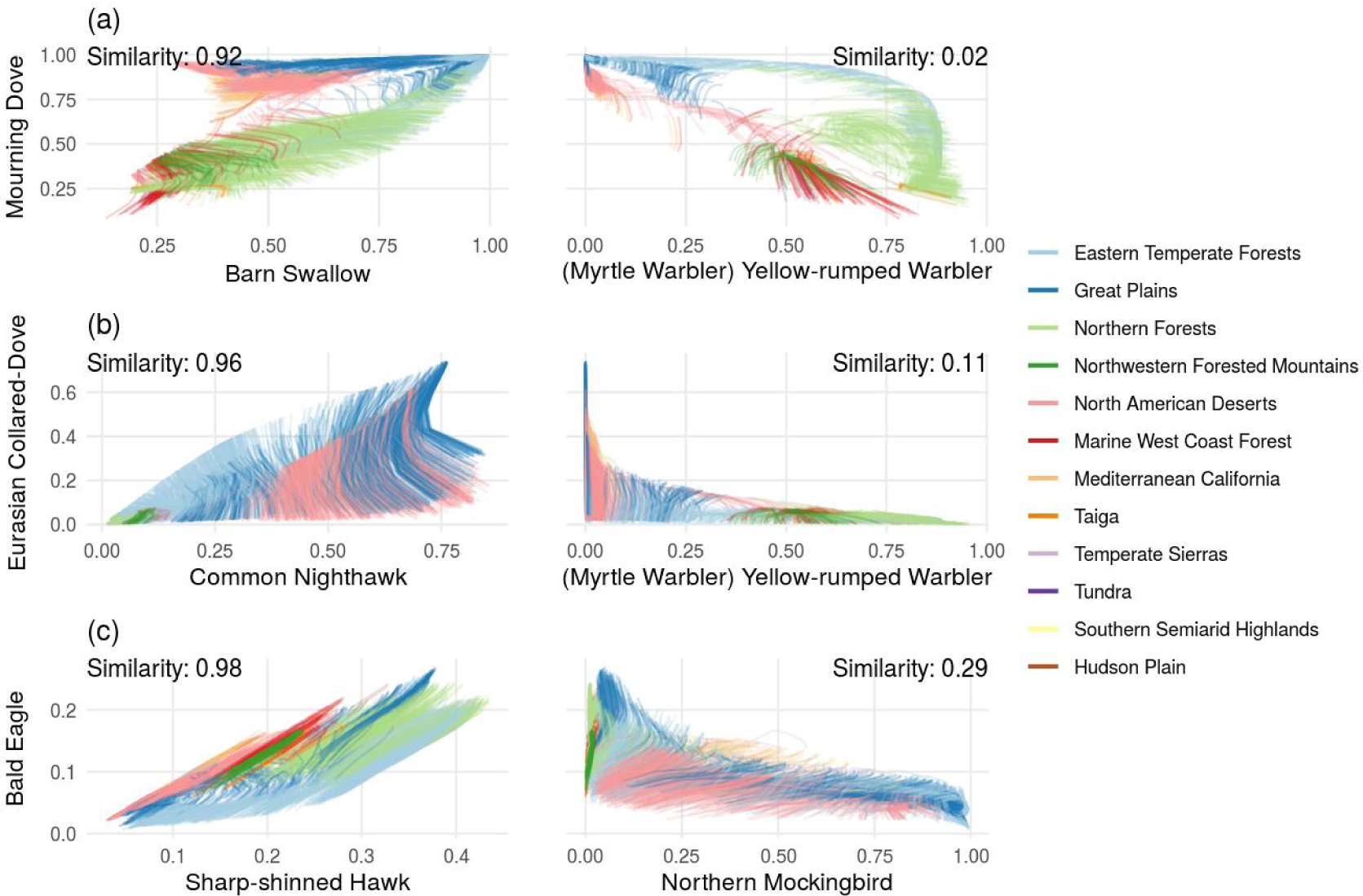
Relationships among species estimated occupancy probabilities through time, by location for species pairs that are nearest neighbors in parameter cosine distance (left column), and most cosine dissimilar (right column). Color indicates EPA level 1 ecoregions. Cosine similarity values for species pairs are printed in the upper corners of each panel. Each route is a line segment that connects estimates of occupancy probabilities for each year from 1997 to 2018. Species pairs with high cosine similarity tend to have positively related occupancy trajectories.

## Discussion

Neural hierarchical models provide a bridge between hierarchical models built from scientific knowledge, and neural networks that approximate functions over structured domains. This framework integrates research in science-based deep learning and ecological modeling, and can use existing hierarchical models as a starting point. The case study provides a proof of concept example of constructing a scalable and performant neural hierarchical model based on a multi-species dynamic occupancy model, providing insights about colonization and extinction dynamics of North American bird assemblages at a continental scale.

The breeding bird survey case study indicates that neural hierarchical models can outperform simpler models. Notably, the final model was performant both quantitatively and qualitatively, detecting population increases and range expansions that are consistent with prior work. The case study extends previous analyses of persistence probabilities at range edges (Royle & Kéry 2007), indicating that common species tend to have higher persistence and colonization probabilities at routes close to their BBS range centroid. Yet, interpreting this result is complicated by irregular range geometry which can lead to centroids that near the edge (or even outside) of a species range, and divergence between actual and estimated ranges due to incomplete sampling (Sagarin & Gaines 2002; Fortin *et al*. 2005; Dallas *et al*. 2017; Knouft 2018). Additional complexity is apparent in Fig. 4c, which indicates that gradients in colonization and persistence may not be isotropic (the same in every direction). With those caveats, the results are broadly consistent with a theoretical expectation that range boundaries can arise from gradients in local extinction and colonization rates (Holt & Keitt 2000).

The case study used a gated recurrent neural network architecture to handle temporal structure (Chung *et al*. 2014), and architectures designed for other data structures present additional opportunities for ecological applications. For example, neural hierarchical models could be used to couple a convolutional neural network observation model for camera trap images (Norouzzadeh *et al*. 2018; Tabak *et al*. 2019) with an ecological process model that describes animal density, movement, or community composition (Burton *et al*. 2015). Gridded data such as modeled climate data, remotely sensed Earth observations, and even 96 well microplates also might use convolutional neural network architectures (Rawat & Wang 2017). Appendix S2 provides a simulated example where gridded data (aerial imagery) are mapped to a state transition matrix of a hidden Markov model. In addition, many ecological datasets exhibit graph structure related to phylogenies, social networks, or network-like spatial structure. While it is possible to adapt convolutional neural networks to operate on distance matrices computed from graphs (Fioravanti *et al*. 2018), graph representation learning can also provide embeddings for nodes that encode network structure and node attributes (Hamilton *et al*. 2017; Cai *et al*. 2018).

Neural hierarchical models can scale to data that are too large to fit in computer memory. Indeed, memory limitations precluded comparison of a fully Bayesian multi-species dynamic occupancy model against the multi-species neural hierarchical model. The current state of the art multi-species occupancy models use approximately one tenth of the number of species, at about half the number of sites, and are static in the sense that colonization and extinction dynamics through time are not represented (Tobler *et al*. 2019). Species distribution models are increasingly scalable due to advances in approximate Gaussian process models (Tikhonov *et al*. 2019), but multi-species dynamic occupancy models have not previously been reported at this scale. This is particularly relevant for extensions of the BBS case study, given the volume of (imperfect) bird data accumulating through citizen science programs (Sullivan *et al*. 2009). The key strategy providing scalability in the case study is stochastic optimization that uses mini-batches - small subsets of a larger dataset - to generate noisy estimates of model performance and partial derivatives that can be used during training (Ruder 2016). With this approach, the entire dataset does not need to be loaded into memory at the same time. This could be useful as a way to scale models that integrate BBS, eBird, and other data (Ngiam *et al*. 2011; Pacifici *et al*. 2017; Zipkin *et al*. 2017; Lin *et al*. 2019).

Though neural hierarchical models can perform well in the large data regime, they might be overparameterized in a small data setting. For this reason, performance comparisons with simpler baseline models are useful. Appendix S2 provides a simulated example, where simple baselines work better with small datasets, and more complex neural hierarchical models work better with large datasets. Approaches familiar to ecologists such as k-fold cross validation and (to a lesser extent) information criteria such as the Akaike Information Criterion (AIC) have been used with neural networks, but such approaches are less common than cross-validation with training, validation, and test partitions of the data (Anderson & Burnham 2004; Jiang & Chen 2016; Ran & Hu 2017). With large datasets, considerable computational resources are required for training, so that retraining the same model many times might make k-fold cross validation infeasible. As a practical matter, even if a neural hierarchical model demonstrates superior predictive performance, a simpler model might be preferred if interpretability and/or explainability is a high priority, as it might be in a decision-making context (Rudin 2018).

Neural networks have a reputation for being “black-box” models. However, the interpretation and explanation of such models is an active area of research (Roscher *et al*. 2019). Interpretations map abstract concepts to domains that humans can make sense of, e.g., mapping neural network parameters to species identity or spatial location (Montavon *et al*. 2018). Explanations are collections of features in such a domain that contributed to a prediction (Montavon *et al*. 2018), e.g., sensitivity of model output to perturbations of the input (Olden & Jackson 2002; Gevrey *et al*. 2003).

Looking ahead, in a forecasting or decision-making setting, it would be important to estimate both aleatoric uncertainty (arising from noise in an observation process) and epistemic uncertainty (arising from uncertainty in a model and its parameters) (Clark *et al*. 2001; Kendall & Gal 2017). Recent advances in accounting for uncertainty in neural network parameter estimates and architectures could be applied in future work. Approaches include “Bayes by backpropagation” (Blundell *et al*. 2015), normalizing flows for variational approximations (Kingma *et al*. 2016), adversarial training (Lakshminarayanan *et al*. 2017), methods that use dropout or its continuous relaxation (Gal *et al*. 2017), and ensemble approaches (McDermott & Wikle 2017). These methods can also help explain neural networks, e.g., to probabilistically estimate sensitivity to model inputs to random masking (Chang *et al*. 2017), or to decompose predictive uncertainty into component parts (Thiagarajan *et al*. 2019).

The potential for combining neural networks with mechanistic models was recognized more than thirty years ago (Psichogios & Ungar 1992; Meade Jr & Fernandez 1994; Lagaris *et al*. 1998). This potential is more easily realized today due to methodological spillover from deep learning into the natural sciences, but also increases in computing power, availability of ecological data, and the proliferation of educational content for quantitative ecology and machine learning. Further, modern deep learning frameworks provide abstractions that allow users to focus on model construction rather than the details of implementation, increasing accessibility in the same way that WinBUGS, JAGS, OpenBUGS, and Stan have done (Lunn *et al*. 2000; Plummer & others 2003; Spiegelhalter *et al*. 2005; Carpenter *et al*. 2017).

Although deep learning and ecological modeling may seem to be separate activities, neural hierarchical models bridge these disciplines. Given the increasing availability of massive ecological data, scalable and flexible science-based models are increasingly needed. Neural hierarchical models satisfy this need, and can provide a framework that links imperfect observational data to ecological processes and mechanisms by construction.

## Acknowledgements

Thanks to David Zonana and Roland Knapp for discussions on the potential of neural hierarchical models, and to Susie Ellis for providing feedback on a draft of the manuscript. This manuscript was also greatly improved by three anonymous reviewers. Also thanks to Brandon Edwards for developing the bbsBayes R package, which was used to acquire and parse the North American Breeding Bird Survey data. This work was motivated by a workshop on machine learning in Earth science, hosted by Earth Lab and organized by the Federation of Earth Science Information Partners and the National Aeronatics and Space Administration Advanced Information Systems Technology program. This work was made possible by the CU Boulder Grand Challenge initiative and the Cooperative Institute for Research in the Environmental Sciences through their investment in Earth Lab. Thanks also to the thousands of U.S. and Canadian participants who annually perform and coordinate the North American Breeding Bird Survey.

## Appendix S1

This appendix includes example specifications for neural occupancy, N-mixture, and hidden Markov models. The goal in providing these specifications is to give concrete examples of neural hierarchical models that are relatively simple, while drawing connections to existing models. PyTorch implementations in Jupyter notebooks are available at https://github.com/mbjoseph/neuralecology.

Generally, the construction of a hierarchical model requires:

1. A **process model** that represents some ecological dynamics or states that can depend on unknown parameters.
2. An **observation model** that relates available data to the process model, possibly dependent on unknown parameters.
3. A **parameter model** for unknown quantities (e.g., their dependence on some input, or relationship to each other). Traditionally, this is discussed in terms of prior distributions, though there may be rich structure encoded in a parameter model as well (Wikle 2003). For a fully Bayesian hierarchical model, all unknowns are represented by probability distributions. In an empirical Bayesian hierarchical model, point estimates are typically used for top-level parameters instead, so that these top-level parameters are treated as fixed, but unknown (Cressie *et al*. 2009). In the examples below, neural networks are applied at the parameter model stage to learn a mapping from inputs to parameters of process and observation models.

Given these components, parameter estimation can proceed in a few ways, depending on the inferential framework used. A loss function can be constructed to find the parameter values that maximize the probability of the data, possibly with some penalties for model complexity (in a maximum likelihood/penalized maximum likelihood framework), find the most probable parameter values, conditional on the data (in a maximum *a posteriori* framework), or compute a probability distribution for all unknowns, conditional on the data (in a Bayesian framework). For many models applied in deep learning, it is often the case that fully Bayesian inference is complicated by high-dimensional multimodal posterior geometry.

The examples here focus primarily on penalized maximum likelihood methods, though Bayesian approaches stand out as a key area for future work (see main text for relevant citations). Often, *L*_2_ norm penalties (also referred to as “weight decay”) are applied on the parameters of neural networks in the loss function to penalize model complexity in an effort to avoid overfitting. Additional strategies include early stopping, where an iterative stochastic optimization scheme is terminated before the training set loss stabilizes (or when the validation set loss begins to increase), which is particularly useful for complex models that can overfit quickly. Finally, unlike maximum likelihood estimation for simple models, because of the multimodality of the likelihood surface for neural network parameters, there are no guarantees that a global maximum exists, or that a particular optimum is not a local maximum. In practice, this is not a major limitation. For more discussion, see Goodfellow *et al*. (2016).

### A single-species single-season neural occupancy model

A single-species single-season occupancy model estimates presence/absence states from imperfect detection/non-detection data (MacKenzie *et al*. 2002). Assume that *n* spatial locations are each surveyed *k* times, during a short time interval for which it is reasonable to assume that the presence or absence of a species is constant. Each spatial location has some continuous covariate value represented by *x*_*i*_ for site *i* = 1, …, *n*, that relates to occupancy and detection probabilities.

#### Observation model

Observations at site *i* consist of *k* surveys, where each survey results in a detection or non-detection. Let *y*_*i*_ represent the number of surveys at site *i* for which a species is detected, and *z*_*i*_ represent the true presence/absence state (if the species is present: *z*_*i*_ = 1; if absent: *z*_*i*_ = 0). Assuming that the probability of detecting a species on a survey conditional on presence is *p*_*i*_, and that each survey is conditionally independent, the observations can be modeled with a Binomial distribution:

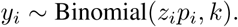

#### Process model

The true presence/absence occupancy state *z* can be treated as a Bernoulli random variable, with occupancy probability *ψ*_*i*_:

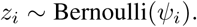

#### Parameter model

A simple approach to account for the effect of the site-level covariate on occupancy and detection would be to include a slope and intercept parameter specific to each component, using the logit function to ensure that estimated probabilities are bounded between 0 and 1:

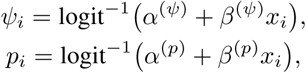

where *α*^*j*^ and *β*^*j*^ are intercept and slope parameters for component *j*.

In contrast, a neural hierarchical model might instead account for site-level variation by modeling occupancy and detection probabilities as outputs of a neural network:

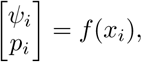

where *f* (*x*_*i*_) is a neural network that takes a scalar as input (the site-level covariate *x*_*i*_) and outputs a two dimensional vector containing occupancy and detection probabilities:

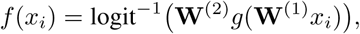

where **W**^(1)^ and **W**^(2)^ are parameter matrices for the first and second layer, *g* is a differentiable nonlinear activation function, e.g., the rectified linear unit activation function (Nair & Hinton 2010), and the inverse logit transform is applied element-wise to ensure that *ψ*_*i*_ and *p*_*i*_ lie between 0 and 1.

#### Loss function

The negative log observed data likelihood can be taken as the loss function (MacKenzie *et al*. 2002). Marginalizing over *z* and treating sites as conditionally independent yields:

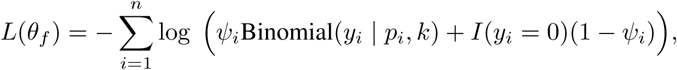

where *θ*_*f*_ represents the parameters of the neural network *f*, and *I*(*y*_*i*_ = 0) is an indicator function equal to one when *y*_*i*_ = 0 (zero otherwise).

The loss for any particular site can be computed efficiently using the log-sum-exp trick, which is particularly useful when the log of summands is available (i.e., 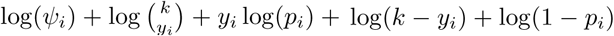 and log(1 − *ψ*_*i*_) are already computed), as is the case in both Stan and PyTorch, which provide the log probability from the Binomial distribution.

#### A single-species neural dynamic occupancy model

A single-species dynamic occupancy model can be used to estimate rates of colonization and extinction when detection is imperfect and sites are repeatedly sampled at multiple time points (MacKenzie *et al*. 2003). Assume that for *T* timesteps, *n* spatial locations are each surveyed *k* times, during a short time interval for which it is reasonable to assume that the presence or absence state is constant. Among timesteps, the true occupancy states of sites can change. Each spatial location has some continuous covariate value represented by *x*_*i*_ for site *i* = 1, …, *n*, that relates to occupancy and detection probabilities.

#### Observation model

Observations at site *i* in timestep *t* consist of *k* surveys, where each survey results in a detection or non-detection. Let *y*_*i,t*_ represent the number of surveys at site *i* in timestep *t* for which a species is detected, and *z*_*i,t*_ represent the true presence/absence state (if the species is present: *z*_*i,t*_ = 1; if absent: *z*_*i,t*_ = 0). Assuming that the probability of detecting a species on a survey conditional on presence is *p*_*i*_, the observations can be modeled using a Binomial distribution:

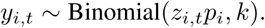

#### Process model

Sites can transition from being unoccupied (*z*_*i,t*_ = 0) to occupied (*z*_*i,t*+1_ = 1) due to colonization, or from being occupied (*z*_*i,t*_ = 1) to unoccupied (*z*_*i,t*+1_ = 0) due to extinction. Let the initial occupancy state be treated as a random Bernoulli variable with probability of occupancy *ψ*_*i*,1_:

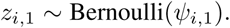

Subsequent occupancy dynamics at site *i* for timesteps *t* = 1, …, *T* are related to the probability of colonization (_*i*_) and the probability of persistence (*ϕ*_*i*_), where the extinction probability is taken to be the complement of persistence (1 − *ϕ*_*i*_).

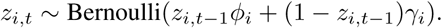

#### Parameter model

In this example, heterogeneity among sites was accounted for using a single layer neural network that ingests the one dimensional covariate *x*_*i*_ for site *i*, passes it through a single hidden layer, and outputs a four dimensional vector of probabilities containing the probabilities of initial occupancy (*ψ*_*i*,1_), persistence (*ϕ*_*i*_), colonization (*γ*_*i*_), and detection (*p*_*i*_):

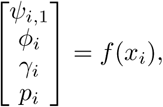

where *f* is a neural network. Concretely, *f* was parameterized as follows:

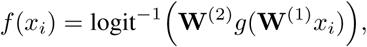

where **W**^(1)^ is a parameter matrix that generates activations from the inputs, *g* is the ReLU activation function, **W**^(2)^ is a parameter matrix that maps the hidden layer to the outputs, and logit^−1^ is the element-wise inverse logistic (sigmoid) function.

#### Loss function

The negative log observed data likelihood was used as the loss function, implemented in PyTorch following the description in MacKenzie *et al*. (2003) (equation 5-6), assuming that detection histories for site *i* = 1, …, *n* are conditionally independent and scaling the forward probabilities to avoid underflow as described in Rabiner (1989).

#### A neural N-mixture model

An N-mixture model can be used to estimate latent integer-valued abundance when unmarked populations are repeatedly surveyed and it is assumed that detection of individuals is imperfect (Royle 2004). Assume that *J* spatial locations are each surveyed *K* times, in a short time interval for which it is reasonable to assume that the number of individuals is constant within locations *j* = 1, …, *J*. Each spatial location has some continuous covariate value represented by *x*_*j*_, that relates to detection probabilities and expected abundance.

#### Observation model

Observations at site *j* in survey *k* yield counts of the number of unique individuals detected, denoted *y*_*j,k*_ for all *j* and all *k*. Assuming that the detection of each individual is conditionally independent, and that each individual is detected with site-specific probability *p*_*j*_, the observations can be modeled with a Binomial distribution where the number of trials is the true (latent) population abundance *n*_*j*_:

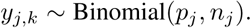

#### Process model

The true population abundance *n*_*j*_ is treated as a Poisson random variable with expected value *λ*_*j*_:

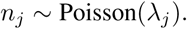

#### Parameter model

Heterogeneity among sites was accounted for using a single layer neural network that ingests the one dimensional covariate *x*_*i*_ for site *i*, passes it through a single hidden layer, and outputs a two dimensional vector containing a detection probability *p*_*i*_ and the expected abundance *λ*_*i*_:

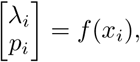

where *f* is a neural network with two dimensional output activations **h**(*x*_*i*_) computed via:

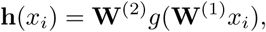

and final outputs computed using the log and logit link functions for expected abundance and detection probability:

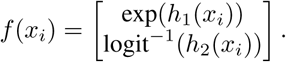

Here too **W**^(1)^ is a parameter matrix that generates activations from the inputs, *g* is the rectified linear unit activation function, and **W**^(2)^ is a parameter matrix that maps the hidden layer to the outputs. Additionally *h*_1_(*x*_*i*_) is the first element of the output activation vector, and *h*_2_(*x*_*i*_) the second element.

#### Loss function

The negative log likelihood was used as the loss function, enumerating over a large range of potential values of the true abundance (from min(*y*_*j*_.) to 5 × max(*y*_*j*_.), where *y*_*j*._ is a vector of counts of length *K*) to approximate the underlying infinite mixture model implied by the Poisson model of abundance (Royle 2004). It is also worth noting that alternative specifications based on a multivariate Poisson model are possible (Dennis *et al*. 2015).

#### A neural hidden Markov model: capture-recapture-recovery

Consider a capture-recapture-recovery study aimed at estimating time-varying parameters (King 2012). Assume that individuals *i* = 1, …, *N* are initially captured, marked, and released on timestep *t* = 0.

The state of an individual *i* on time step *t* is denoted *z*_*i,t*_. Individuals are either alive (*z*_*i,t*_ = 0), recently dead such that their bodies are discoverable (and marks identifiable) (*z*_*i,t*_ = 1), or long dead such that their bodies are not discoverable and/or marks are no longer identifiable (*z*_*i,t*_ = 2).

#### Observation model

Observations *y*_*i,t*_ are made on time steps *t* = 1, …, *T*, and if an individual is detected alive on timestep *t, y*_*i,t*_ = 1, and if an individual is detected as recently dead *y*_*i,t*_ = 2, otherwise if the individual is not detected *y*_*i,t*_ = 0.

Let **Ω**_*t*_ denote a time-varying emission matrix containing state-dependent observation probabilities:

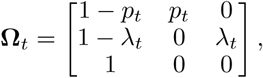

where *p*_*t*_ is the probability of detecting an individual on timestep *t*, conditional on the individual being alive, and *λ*_*t*_ is the probability of recovering a recently dead individual on timestep *t* that has died since timestep *t* − 1. The rows of the emission matrix correspond to states (alive, recently dead, long dead), and the columns correspond to observations (not detected, detected alive, detected dead).

#### Process model

At time *t*, the transition probability matrix **Γ**_*t*_ contains the probability of transitioning from row *j* to column *k*:

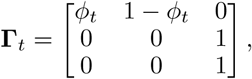

where survival probability is *ϕ*_*t*_ = *P* (*z*_*i,t*+1_ = 0 | *z*_*i,t*_ = 0), dead individuals stay dead, and the rows and columns of the transition matrix correspond to states (alive, recently dead, long dead).

#### Parameter Model

Heterogeneity among timesteps was accounted for using three two-layer neural networks (one for *ϕ*, one for *λ*, and one for *p*). Each network ingests a univariate time series and outputs a corresponding time series of parameter values (i.e., each network maps a sequence of inputs *x* = (*x*_*t*=1_, …, *x*_*t*=*T*_)′ to a sequence of parameter values, e.g., *p* = (*p*_*t*=1_, …, *p*_*t*=*T*_)′ for the detection probability network:

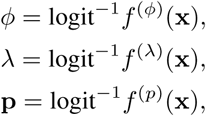

where *f* ^(*j*)^ is a neural network for parameter *j*.

#### Loss function

The negative log observed data likelihood was used as the loss function, and computed using the forward algorithm (Zucchini *et al*. 2017). Observation histories for individuals were assumed to be conditionally independent. As an aside, one can specify such models in terms of the complete data likelihood (i.e., in terms of the hidden states) using programming frameworks that implement automatic enumeration of discrete latent variables with finite support, such as Pyro (Bingham *et al*. 2018).

## Appendix S2

This appendix describes an animal movement model parameterized by a convolutional neural network, including simulations to explore how the amount of training data affects performance relative to simpler baseline models. Convolutional neural networks (CNNs) are widely used in computer vision applications, acting as function approximators that take an image as input and output a vector of probabilities that the image is a picture of a cat, dog, car, etc. (Rawat & Wang 2017). In ecological contexts, CNNs have been used to identify plants and animals (Norouzzadeh *et al*. 2018; Fricker *et al*. 2019; Tabak *et al*. 2019). However, CNNs considered more broadly as function approximators have additional uses. For example, instead of mapping an image to a vector of class probabilities, a CNN might map an image to a state transition probability matrix in a hidden Markov model, such as those used for animal movement trajectories (Zucchini *et al*. 2017).

Generally, the movement of an individual animal depends on the spatiotemporal context around their location at any time, and on their behavioral state. In many animal movement models, covariates for the state transitions are included using a linear combination of values followed by a softmax transformation to ensure probabilities within rows of a state transition probability matrix sum to one. This allows inference about spatiotemporal covariates that explain movement such as distance to water, temperature, hour of day, wind, and ocean surface current (Patterson *et al*. 2017; Johnson *et al*. 2018; McClintock & Michelot 2018a). In most cases, only the values of the covariates at point locations get used. However, it is reasonable to expect that the spatiotemporal context around these point locations could also be relevant.

Instead of extracting values at points, consider using gridded raster data around point locations such as image “chips” centered around observed point locations containing contextual data within some spatiotemporal window. Input rasters might contain satellite and/or aerial imagery, continuous or categorical landscape features (e.g., a digital elevation model or land cover map), or meteorological data (e.g., gridded temperature or wind speed data).

### Scenario description

Consider an animal that prefers to forage in and around tree canopies, but frequently moves between canopies over bare ground. To generate a semi-realistic simulation of this scenario, movement trajectories were simulated over vegetation canopy height models from the San Joaquin Experimental Reserve, one of the core National Ecological Observatory Network (NEON) sites (Keller *et al*. 2008). These canopy height models are estimated using lidar data on the NEON Airborne Observation Platform (AOP) - a plane that flies over NEON sites regularly and collects a variety of data, including high resolution orthorectified aerial imagery (Fig. S2.1).

In this simulation, the “correct” covariate data that affects state transitions – which is rarely known in real systems – is the height of the tree canopy. Assume that canopy height is unknown, but aerial imagery (or high resolution satellite imagery) is available. This aerial red-green-blue (RGB) imagery should to relate in some way to canopy height, but the mapping from an RGB image to canopy height is likely complex. This provides an opportunity to test whether a convolutional neural hierarchical model can learn such a mapping using information about animal movement.

#### Simulating animal movement through tree canopies

Individual animal movement trajectories were simulated over a real canopy height model of the San Joaquin Experimental Reserve. The spatial region of interest is a 6 km by 5 km region, within which a 2018 NEON AOP mission was flown that generated a coregistered 1 m canopy height model (product code DP3.30015.001), and 10 cm high resolution orthorectified RGB camera reflectance data (product code DP3.30010.001). Due to some extreme outliers in the canopy height model, all values greater than 30m (∼0.0025% of cells) were set to 30m. Then, the resulting raster was max-scaled (dividing by the maximum) to compress the range of values to the interval [0, 1].

An animal movement model with two states (foraging and in transit) was used to simulate trajectory data. Animals were more likely to forage where the canopy is high, and more likely to be in transit where canopy height is low.

**Figure S2.1:**
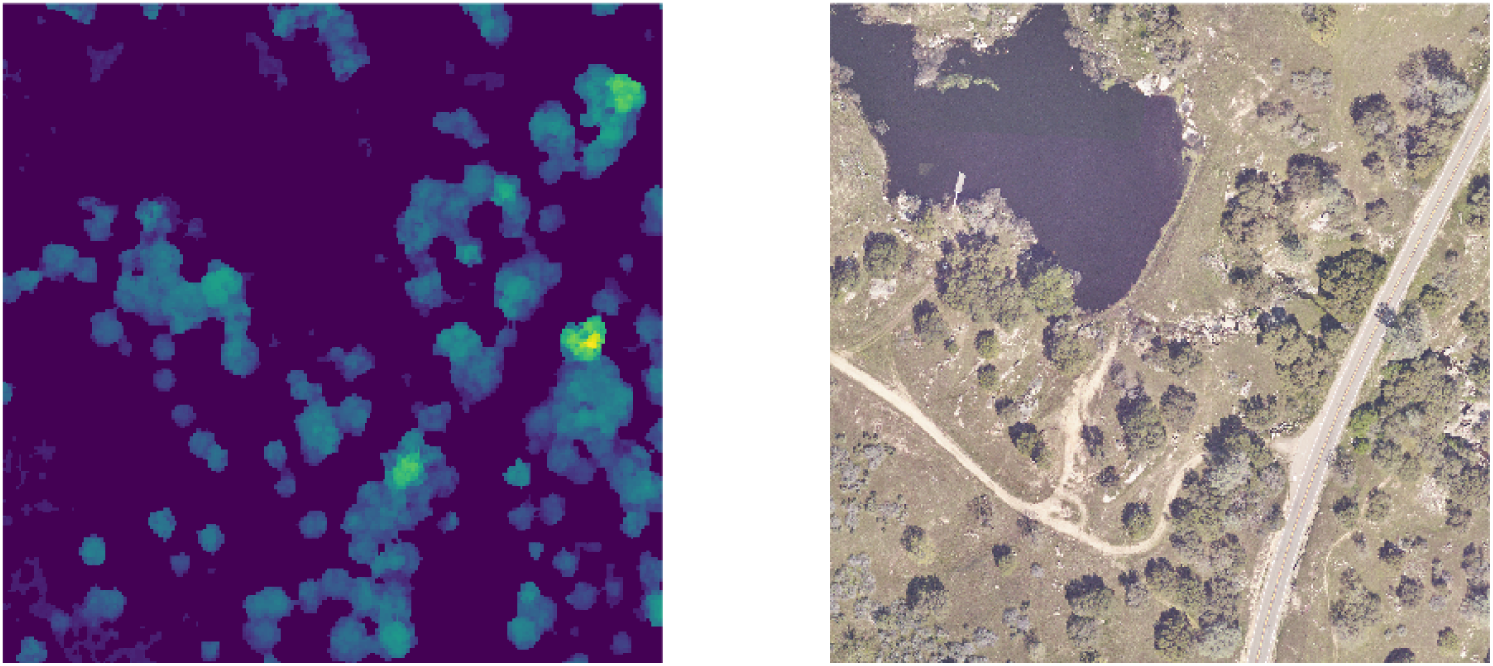
An example of a 1 m pixel resolution canopy height model (left) and corresponding high resolution (10 cm) orthorectified camera imagery over the San Joaquin Experimental Reserve. In the simulation, canopy height affects behavioral state transitions and subsequent animal movement, but the available data might consist only of aerial imagery. A 250 m by 250 m subset of the study area is displayed.

#### Behavioral states

Formally, consider a time series of length *T* containing the state of an animal at discrete times *t* = 1, …, *T*, where *s*_*t*_ = 1 means the animal is “in transit” and *s*_*t*_ = 2 means the animal is “foraging” at time *t*. The state *s*_*t*_ is either 1 or 2 for any particular *t*. The probabilities of transitioning between states is time-varying, and is summarized in a matrix **Γ**^(*t*)^, which contains the transition probabilities 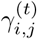 for states *i, j* = 1, 2 at time *t*. Each of these elements provides the probability of transitioning from one state to another, so that 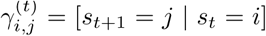. For example 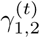 would provide the probability of transitioning from “in transit” in time *t* (*s*_*t*_ = 1) to “foraging” in time *t* + 1 (*s*_*t*+1_ = 2). At the first timestep, the state probabilities are contained in a row vector *δ*, where *δ*_i_ = [*s* _*t*=1_ = *i*], for states *i* = 1 and *i* = 2. In the simulation, the stationary state probabilities at randomly initialized starting locations were used as initial state probabilities.

To ensure that “foraging” was the more likely state in the canopy, and “in transit” was the most likely state over bare ground, the true state transition probabilities in the simulation were modeled as a logit-linear function of canopy height:

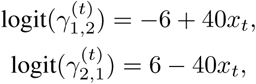

where *x*_*t*_ is the scaled canopy height at an animal’s location in time *t* (Fig. S2.2A). This fully specifies the state transition matrix, because the rows must sum to one, implying for example that 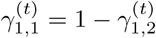.

#### State-dependent movement

The “foraging” and “in transit” behavioral states are associated with different movement patterns. Foraging is characterized by small step lengths with undirected turns. Movement trajectories for animals in transit are characterized by longer step lengths and more directed movements.

**Figure S2.2:**
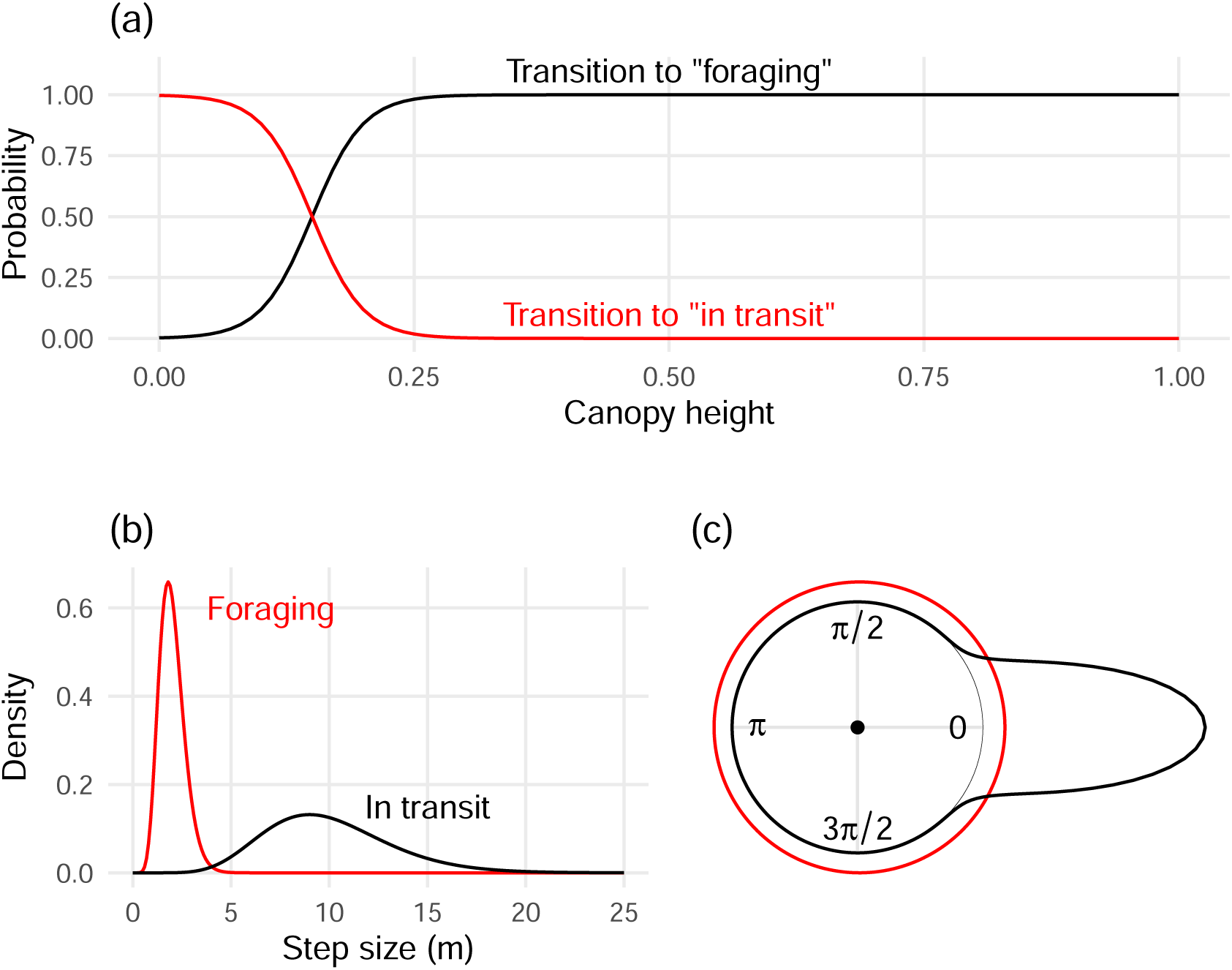
Panel A shows the true relationship between scaled canopy height (x-axis) and state transition probabilities in the simulation. Panels B and C show the densities of step sizes (B) and turn angles in radians (C) used in the simulation, colored by behavioral state.

Formally, if the vector **z**_*t*_ summarizes movement in the interval from time *t* to *t* + 1, it is common to consider two quantities: the step size *l*_*t*_ and turning angle *ϕ*_*t*_, so that **z**_*t*_ = (*l*_*t*_, *ϕ*_*t*_) (Patterson *et al*. 2017). Thus, the movement model is a hidden Markov model with states 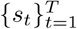, transition probability matrices 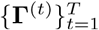, and emissions 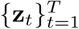.

In the simulation, step sizes were drawn from a gamma distribution with state-dependent parameters (Fig. S2.2A):

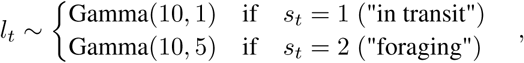

for *t* = 1, …, *T*. Turn angles were drawn from von Mises distributions with state-dependent parameters (Fig. S2.2A):

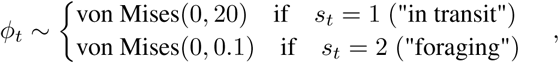

for *t* = 2, …, *T*. Initial movement directions (at *t* = 1) were randomly drawn from the uniform circular distribution, though strictly speaking these are not turn angles which require three points to compute.

**Figure S2.3:**
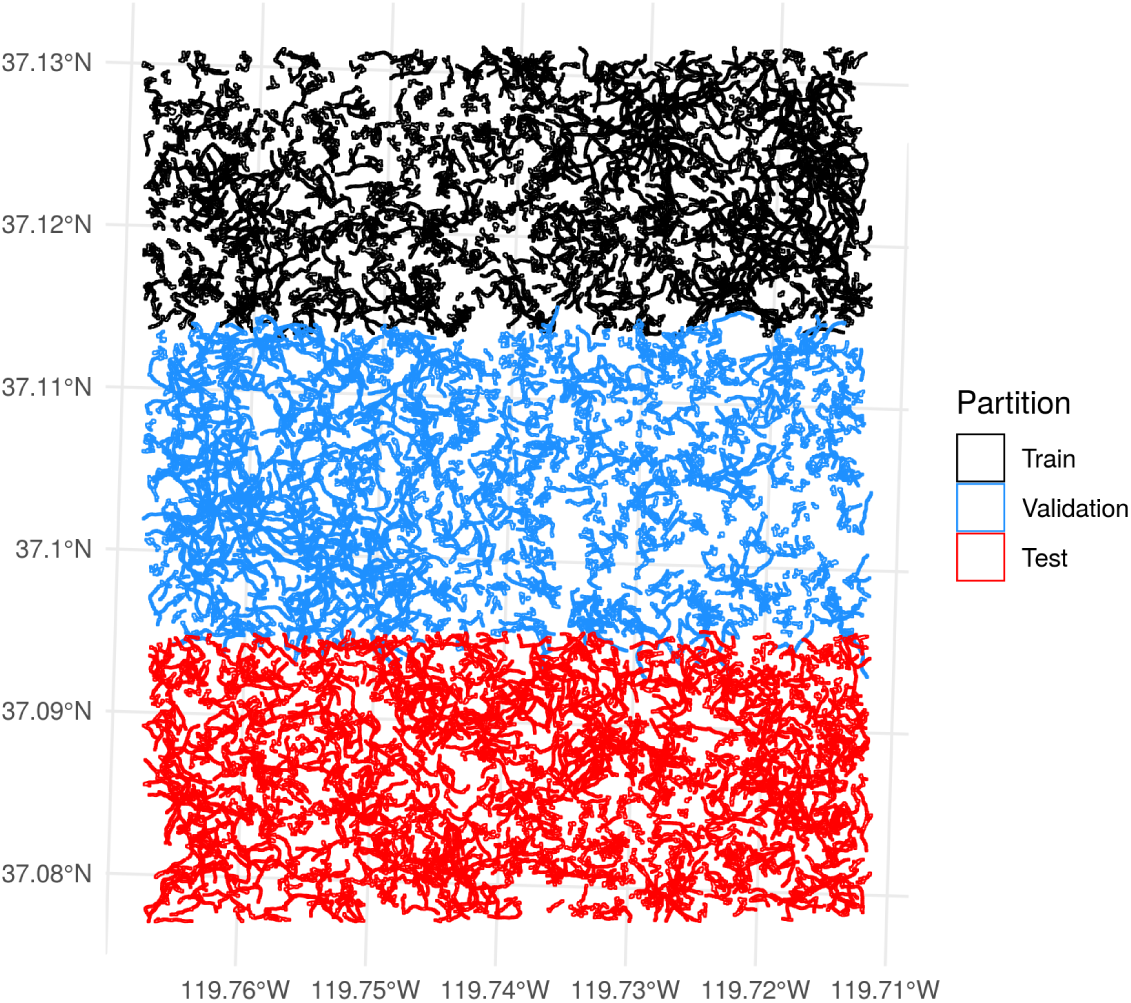
Map of the simulated trajectories in space, colored by dataset partition.

Trajectories simulated within the study area were partitioned into three sets based on the northing coordinate boundaries of the trajectory extents. Trajectories in the northern third of the study area were assigned to the training set, those in in the southern third were used as a withheld test set, and those in the middle third were used as a validation set (Fig. S2.3). For each partition, 1024 trajectories were simulated, to generate 3072 total trajectories across all partitions.

#### Model descriptions

Three models were developed, each of which maps a different set of inputs to transition probability matrices:

1. A **best case model** that takes canopy height as a state transition covariate. This represents the perfect scenario (unlikely in practice) where all relevant spatiotemporal information is provided to the model, and the generative model is correctly specified in every way. This best case model provides a useful upper bound on predictive performance.
2. A **point extraction model** that takes the RGB reflectance values from the aerial imagery extracted at point locations. This represents a more common scenario where covariate data indirectly related to the relevant spatiotemporal information (canopy height) are extracted at point locations. This model is likely to perform poorly, as the RGB reflectance at a point location may not contain much information about canopy height.
3. A **convolutional hidden Markov model** that takes image chips centered on point locations as input, and maps these image chips to transition matrices using a convolutional neural network. If RGB image chips contain more information about canopy height than simple RGB point extractions and sufficient training data are available, this model should perform better than the point extraction model.

**Best case model**

The best case model is the generative model for simulated trajectories. The relationship between canopy height and state transition probabilities is logit linear (as in the simulation), and intercept and slope parameters are estimated:

**Figure S2.4:**
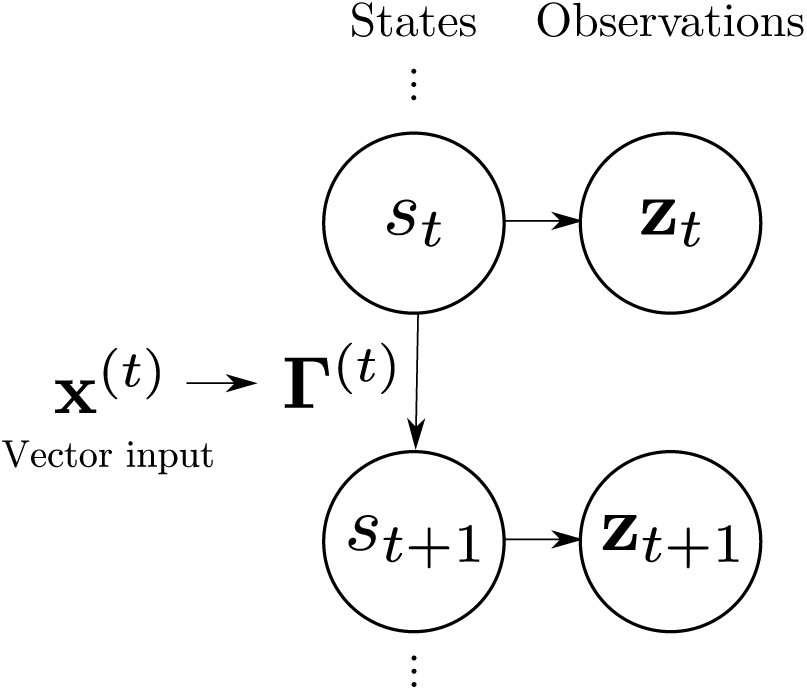
Graphical representation of a generic hidden Markov models for animal movement. The behavioral state *s*_*t*_ is associated with a state transition probability matrix **Γ**^(*t*)^, with observations **z**_*t*_ that represent the movement trajectory of the animal. Inputs contained in a vector **x**^(*t*)^ are mapped to the transition probability matrix by a function (usually a linear combination on a transformed scale).

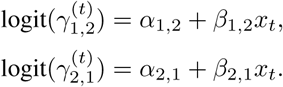

Here as before 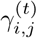 is the probability of transitioning from state *s*_*t*_ = *i* to *s*_*t*+1_ = *j*. Intercept terms are represented by *α*_*i,j*_, and slopes by *β*_*i,j*_, with *x*_*t*_ representing scaled canopy height. This model uses covariates for transition probabilities in the same way that many do: using a linear combination on a transformed scale (Fig. S2.4).

#### Point extraction model

The point extraction model includes parameters to map the RGB image reflectance values (*r*_*t*_, *g*_*t*_, and *b*_*t*_) to the transition probabilities using a linear combination on the logit scale:

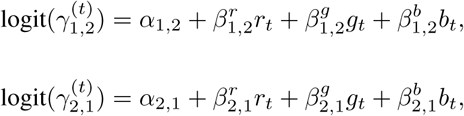

where 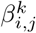 is a coefficient for the (*i, j*)^*th*^ transition probability and image band *k*. This model is also representative of the common approach taken to include covariates in such models - via a linear combination on a transformed scale (Fig. S2.4),

#### Convolutional hidden Markov model

The convolutional hidden Markov model is a neural hierarchical model that maps an image chip **X**^(*t*)^ centered on an animal’s location at time *t* to a transition probability matrix **Γ**^(*t*)^. This is a departure from the previous two models. The input **X**^(*t*)^ is a multidimensional array instead of a real number (in the best case model) or a numeric vector (containing RGB reflectances in the point extraction model).

To generate image chip input arrays, square crops from the aerial RGB imagery were created centered on each simulated location. The spatial footprint of each chip was 128 × 128 pixels (≈ 13 m × 13 m). This created a 3 × 128 × 128 array for each point location along each trajectory, where the three channels correspond to reflectance values in the red, green, and blue spectral bands (Fig. S2.5).

**Figure S2.5:**
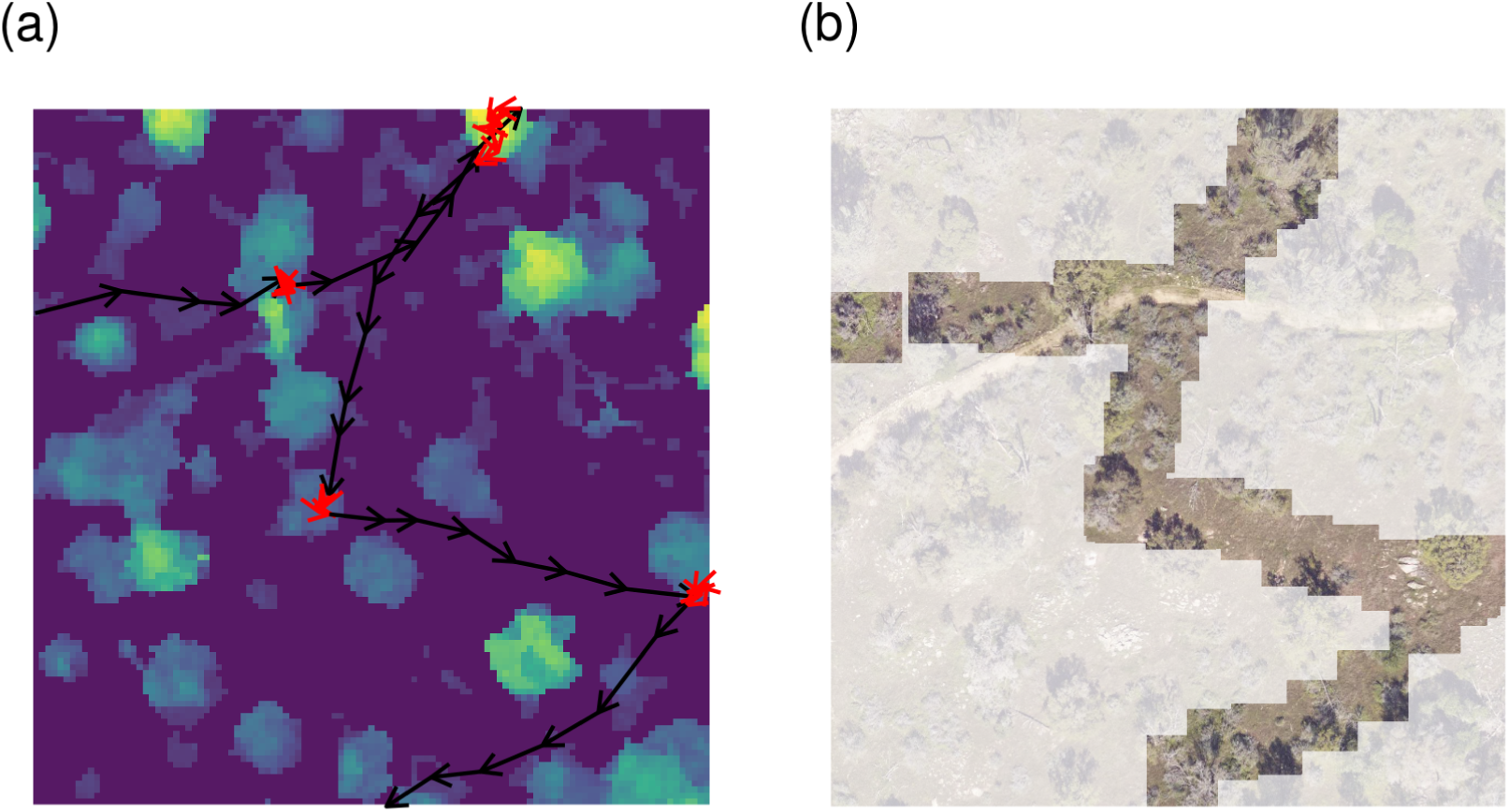
An example of a simulated movement trajectory. Panel (a) shows the true states and scaled canopy height, which determines the state transition probabilities. Red indicates that the animal is foraging, black indicates that it is in transit. The background color map shows the scaled canopy height. Panel (b) shows 128 by 128 image chips centered on point locations in the high resolution aerial imagery, which are used as inputs in the convolutional hidden Markov model of animal movement. Camera imagery outside of these image chips is shown in a lighter shade.

To illustrate the potential for including additional raster data in addition to imagery, an additional band was concatenated to each chip which contained zeros everywhere except for a 2 × 2 region in the center of the 128 × 128 grid, generating 4 × 128 × 128 arrays. In real applications, this might represent additional raster data relevant to movement – inputs need not be images per se.

The convolutional hidden Markov model for animal movement mapped these 4 × 128 × 128 image chips to 2 × 2 state transition probability matrices 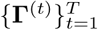 (Fig. S2.6). The architecture of the convolutional neural network is a simplified version of the AlexNet model that plays an important role in the history of deep learning in computer vision (Krizhevsky 2014), though more modern architectures might perform better. Briefly, the input image is passed through a series of 2-dimensional convolutions, followed by nonlinear activation functions, followed by 2d max-pooling layers, creating a 64 × 2 × 2 lower spatial resolution array with many “channels”. This three dimensional array is flattened to a one dimensional array, creating a vector of length 64 × 2 × 2, which is then passed to a series of fully connected hidden layers with nonlinear activations and dropout regularization (Srivastava *et al*. 2014) to create a vector of length 4. This vector is reshaped to a 2 × 2 matrix, then a softmax transformation is applied row-wise to ensure that the row probabilities sum to one (as they should in a state transition probability matrix). The resulting 2 × 2 matrix is the transition probability matrix **Γ**^(*t*)^, generated from the input **X**^(*t*)^.

The precise structure of the convolutional neural network that maps the input raster to the state transition probability matrix was as follows (in PyTorch-like psuedocode):

~~~
Sequential(
  Conv2d(4, 16, kernel_size**=**9, stride**=**3),
  LeakyReLU(),
~~~

**Figure S2.6:**
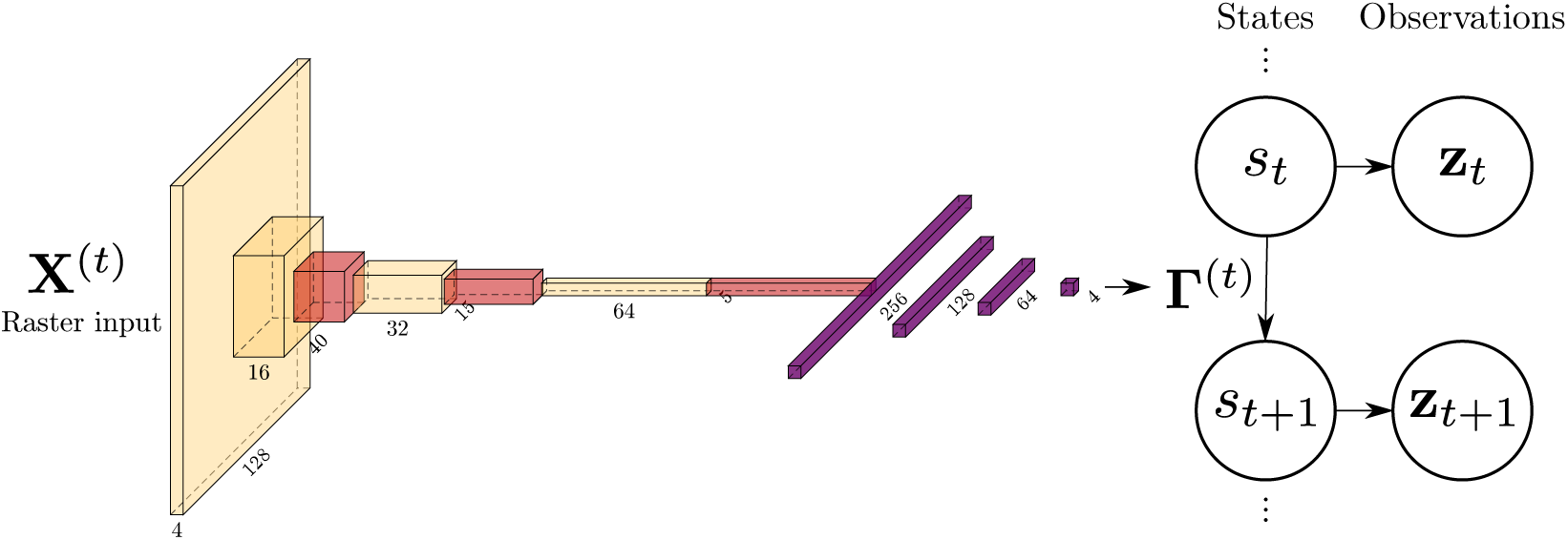
A convolutional neural network that maps a raster (in this case a 4 × 128 × 128 grid) to a state transition probability matrix of a hidden Markov model. Yellow boxes indicate input arrays and outputs from convolutional layers, with labeled dimensions. Red boxes represent two dimensional maximum pooling layers. Purple boxes represent fully connected hidden layers. The final vector of length 4 is reshaped to a 2 by 2 matrix, and then a softmax transform is applied row-wise to ensure that the rows sum to one for the transition probability matrix at timestep *t*, denoted **Γ**^(*t*)^.

~~~
  MaxPool2d(kernel_size**=**3, stride**=**2),
  Conv2d(16, 32, kernel_size**=**5),
  LeakyReLU(),
  MaxPool2d(kernel_size**=**3, stride**=**2),
  Conv2d(32, 64, kernel_size**=**3),
  LeakyReLU(),
  MaxPool2d(kernel_size**=**3, stride**=**2)
  view(**-**1), *# flatten into a vector*
  Dropout(),
  Linear(64 **∗** 2 **∗** 2, 128),
  LeakyReLU(),
  Dropout(),
  Linear(128, 64),
  LeakyReLU(),
  Dropout(),
  Linear(64, 4),
  view(2, 2),
  softmax(dim**=-**1)
)
~~~

In addition to using dropout to regularize the convolutional model, an *L*_2_ penalty of 10^−5^ was also applied. To further regularize the model, image augmentation (random horizontal and vertical flips) was also applied while training, though this might not be desirable in cases where directional orientation of the input image is important (Simonyan & Zisserman 2014). Image augmentation generally includes strategies to perturb data while training computer vision models in an attempt to generate a robust model (i.e., one that is insensitive to translation, orientation, hue, contrast, etc.).

In contrast to more common applications of convolutional neural networks where individual images are labeled (e.g., with bounding boxes and species identities in camera trap imagery), the observed movement trajectories (step sizes and turning angles) can be thought of as analogous to implicit “labels” for this convolutional hidden Markov model.

#### Model comparisons

The predictive performance of the three models (best case, point extraction, and convolutional) was compared across a gradient of training set sizes. This gradient included datasets that consisted of 16, 32, 64, 128, 256, 512, and 1024 simulated trajectories, each of which consisted of 50 timesteps.

**Figure S2.7:**
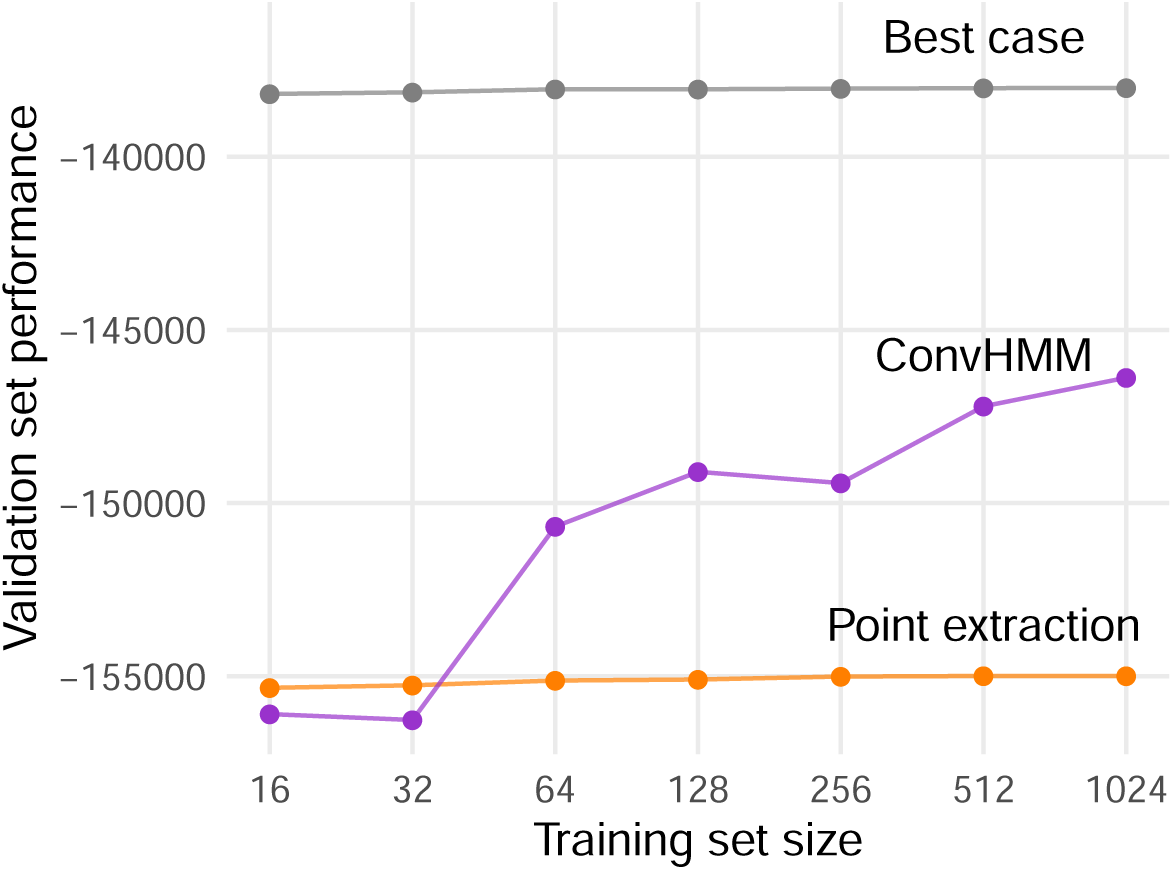
Model performance on withheld validation data. The x-axis is the number of movement trajectories in the training data. The y-axis shows the performance (predictive log-likelihood) on withheld validation data. Each point is the result of a simulation, and colored lines connect results for each model. ConvHMM is the convolutional hidden Markov model of animal movement.

The smaller datasets were subsets of the larger datasets. For each model/training set combination, predictive performance was evaluated using the out of sample log-likelihood, computed for the 1024 withheld validation trajectories using the forward algorithm (Patterson *et al*. 2017). After using the validation data to compare models, the preferred model was retrained using the training and validation data, and final predictive performance was evaluated on the still withheld test set.

### Results

As the training set increased in size to more than 64 trajectories, the convolutional movement model’s performance exceeded the point extraction baseline (Fig. S2.7). When the training set was relatively small, the convolutional movement model performed worse than the point extraction baseline. Validation set performance continued to increase as training data were added for all modles. The rate of increase was greatest for the convolutional model, however, and validation set performance did not appear to saturate even for the largest training data set (Fig. S2.7). As expected, the best case model, which is correctly specified and has access to the “correct” input data (canopy height) always had the highest performance. But, restricting attention to models that only have access to the RGB imagery, the convolutional movement model was preferred.

Final predictive checks on the withheld test set indicated that the convolutional model was able to estimate the true state transition probabilities fairly well. Transition probabilities from “in transit” (*s*_*t*_ = 1) to “foraging” (*s*_*t*_ = 2) were not captured as well as transitions from “foraging” (*s*_*t*_ = 2) to “in transit” (*s*_*t*_ = 1) (Fig. S2.8). In particular, the estimated distribution of transition probabilities was bimodal, but not as sharply peaked as the distribution of true transition probabilities.

Last, to provide a qualitative sense of what the final model had learned, test set image chips with the highest predicted transition probabilities are visualized in Fig. S2.9. Consistent with the underlying generative model, image chips centered on tree canopies were associated with the highest probabilities of transitioning into the “foraging” state, and image chips centered on bare ground (often with intermittent rocks) were associated with the highest probabilities of transitioning into the “in transit” state.

Taken together, these results indicate that simpler models might perform better when limited training data are available, but that neural hierchical models might provide predictive performance improvements for large datasets.

**Figure S2.8:**
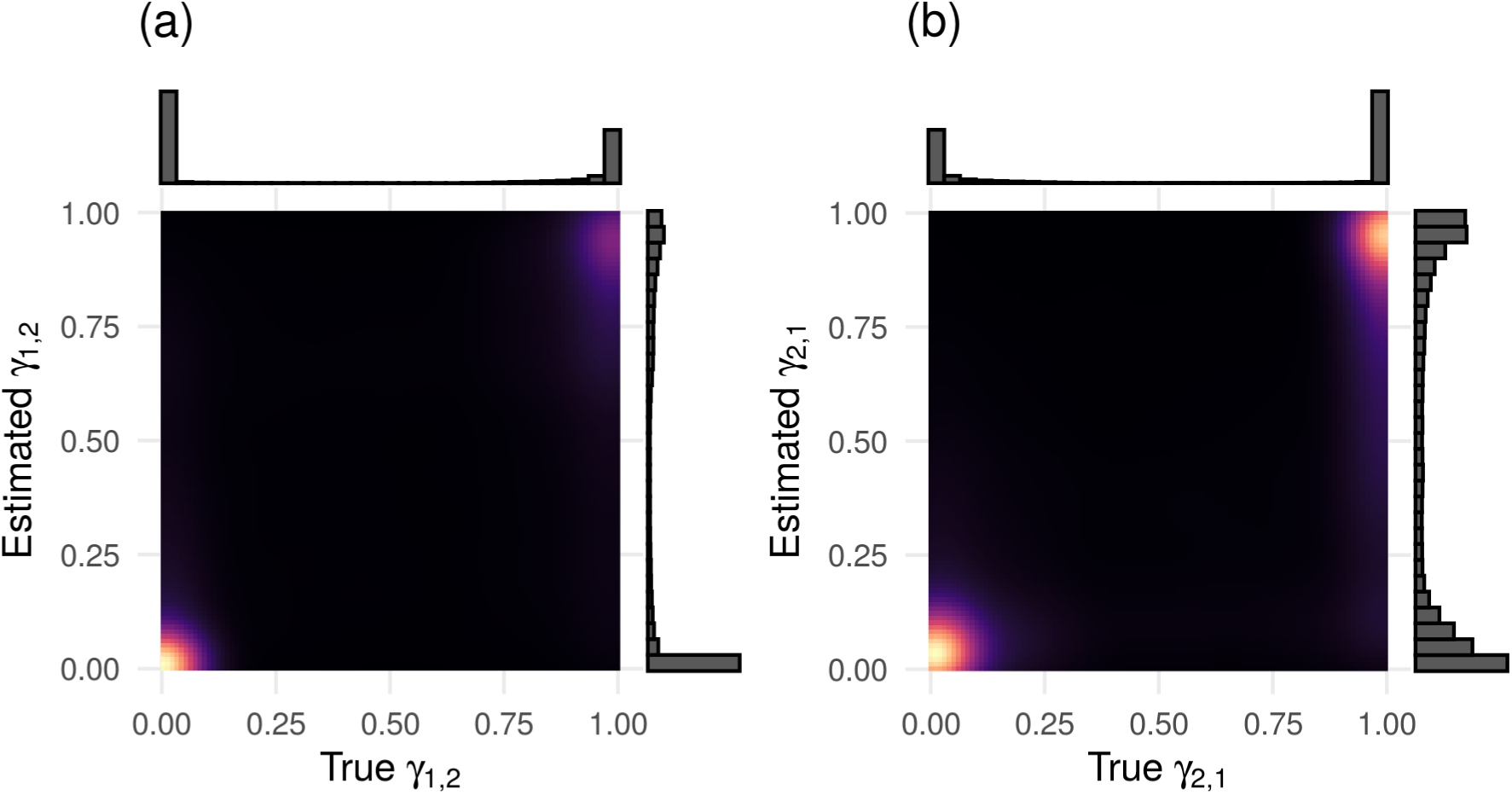
Joint densities of true (x-axis) and estimated (y-axis) state transition probabilities in the final convolutional movement model for the withheld test set. Panel (a) shows the distribution for *γ*_1,2_ (transitions to “foraging”), with marginal histograms for the true and estimated probabilities. Panel (b) shows the same for *γ*_2,1_ (transitions to “in transit”). Cell color represents the density of observations, with brighter colors indicating higher densities.

**Figure S2.9:**
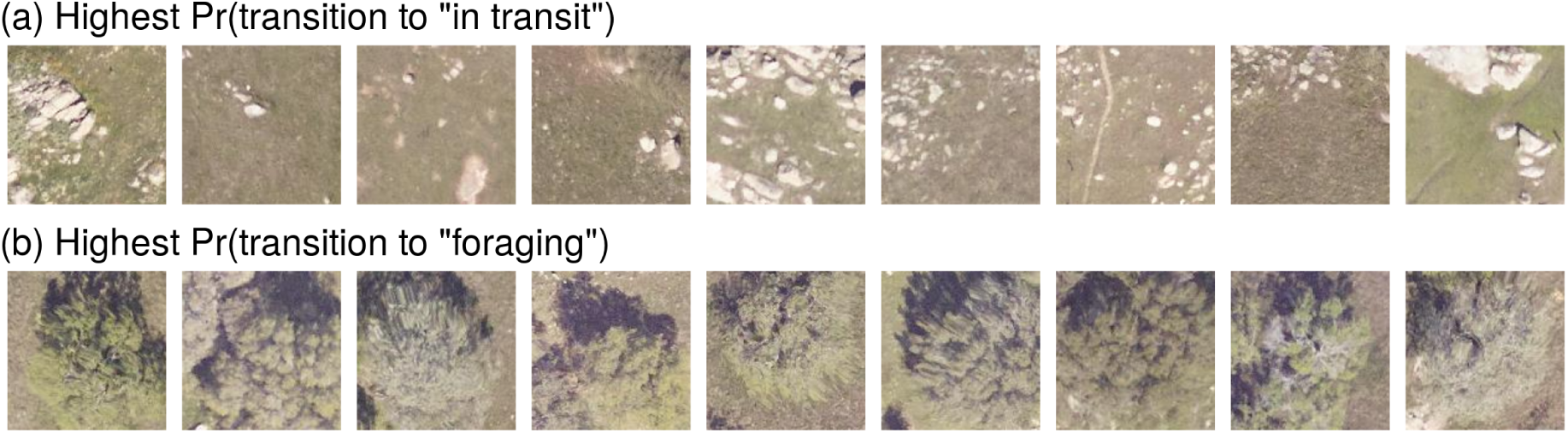
Top nine test set image chips with the highest probabilities of transitioning (a) from a “foraging” state to “in transit", and (b) from “in transit” to “foraging”.

#### Implementation notes

The best case and point extraction models were both fit using the momentuHMM R package (McClintock & Michelot 2018b). The convolutional movement model was implemented with PyTorch (Paszke *et al*. 2017). All code required to reproduce the analysis is available on GitHub at https://www.github.com/mbjoseph/neuralecology.

## Appendix S3

This appendix includes details on the structure and implementation of the baseline and neural dynamic occupancy models for breeding bird survey data.

### Baseline model structure

#### Process model

The baseline model was a hierarchical Bayesian dynamic occupancy model, fit separately to each species. Let *s* = 1, …, *S* index survey routes, *t* = 1, …, *T* index years, and *j* = 1, …, *J* index bird species. Dropping subscripts for species *j*, for any particular species’ model, presence/absence states are characterized by the following dynamics:

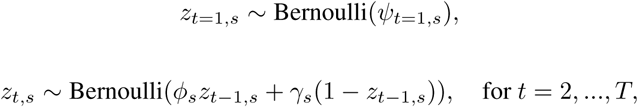

where *ψ*_*t,s*_ is the probability of occurrence at route *s* in year *t, ϕ*_*s*_ is the probability of persistence conditional on presence in the previous year, and *γ*_*s*_ is the probability of colonization conditional on absence in the previous year.

#### Observation model

Detection/non-detection data arise via a binomial observation process:

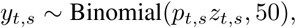

for all *s* and *t*, where the binomial sample size 50 arises from having 50 stops along each route.

#### Parameter model

Heterogeneity in occupancy dynamics was introduced in the model via additive adjustments for EPA level one ecoregions and route-level characteristics (centered and scaled to have mean zero and unit variance):

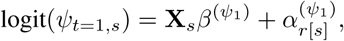

where **X**_*s*_ is row *s* from the design matrix **X** containing the route-level features, 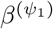 is a parameter vector associated with initial occupancy, and EPA level one adjustments are denoted 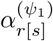, where *r*[*s*] represents the region *r* containing route *s*. Similarly, persistence and colonization probabilities are modeled as:

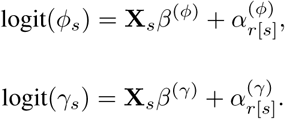

Heterogeneity in detection probability was included as:

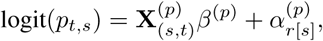

where 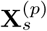 is a augmented version of **X**_*s*_ that adds survey-level features related to detection probability (survey duration, along with start and end sky, temperature, and wind conditions, which vary by route and year), *β*^(*p*)^ is a coefficient vector, and 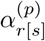 is an ecoregion adjustment.

#### Prior distributions

Prior distributions were constructed to facilitate borrowing of information across the four parameter dimensions of initial occupancy, persistence, colonization, and detection. Ecoregion adjustments were assigned multivariate normal priors:

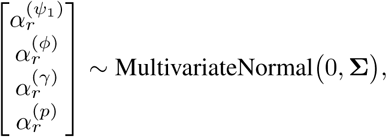

for regions *r* = 1, …, *R*. The covariance matrix **Σ** was constructed for each level as **Σ** = diag(*σ*)**Ω**diag(*σ*), where *σ* is a vector of length 4, diag(*σ*) is a 4 × 4 diagonal matrix with entries equal to *σ*, and **Ω** is a 4 × 4 correlation matrix.

Prior distributions were specified as follows:

*σ* ∼ Gamma(1.5, 10),

**Ω** ∼ LKJ-Correlation(10),

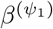 ∼ Normal(0, 1),

*β*^(*ϕ*)^ ∼ Normal(0, 1),

*β*^(*γ*)^ ∼ Normal(0, 1),

*β*^(*p*)^ ∼ Normal(0, 1).

#### Posterior distribution

The posterior distribution is proportional to:

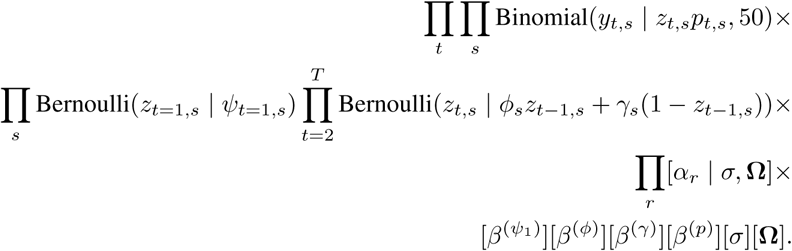

#### Single species neural hierarchical model structure

The single species neural hierarchical model is a dynamic occupancy model parameterized by a neural network.

#### Process model

Presence/absence states are characterized by the following dynamics:

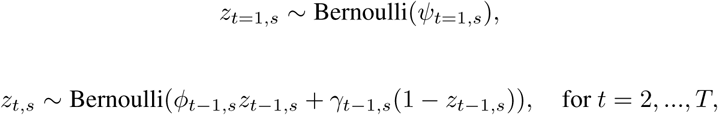

where *ψ*_*t,s*_ is the probability of occurrence at route *s* in year *t, ϕ*_*t*−1,*s*_ is the probability of persistence conditional on presence in the previous year: Pr(*z*_*t,s*_ = 1|*z*_*t*−1,*s*_ = 1) = *ϕ*_*t*−1,*s*_, and *γ*_*t*−1,*s*_ is the probability of colonization conditional on absence in the previous year: Pr(*z*_*t,s*_ = 1|*z*_*t*−1,*s*_ = 0) = *γ*_*t*−1,*s*_.

#### Observation model

Detection data are modeled using a binomial distribution:

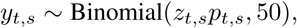

for all *s*, and *t*.

#### Parameter model

Heterogeneity in occupancy and observation parameters was included by modeling initial occupancy, persistence, colonization, and detection probabilities as outputs of a neural network (Fig. S3.1). Categorical inputs to the network included EPA level one ecoregions, which were mapped to 8 dimensional numeric vector embeddings, 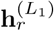 for regions *r* = 1, …, *R* (Guo & Berkhahn 2016). Categorical entity embeddings are equivalent to using one layer of a neural network with 8 dimensional output, where the inputs are one-hot encoded ecoregion codes, but in practice using an embedding layer is more computationally efficient because it capitalizes on the sparsity of a one-hot encoding.

For example, let *R* represent the number of level 1 ecoregions containing BBS routes (*R* = 14). Let 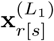 represent a one-hot encoded vector of length *R*, where all entries are zero except for the index of the level 1 ecoregion containing route *s* (denoted *r*[*s*]), which is set equal to one. Then, let 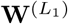. represent a parameter matrix of size 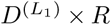, where 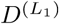 is the dimensionality of the embedding 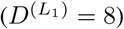. Note that one-hot encodings are often used linear models, where categorical covariates are coded as dummy variables, and this is a special case where the embeddings are one dimensional. Then, the embedding 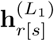 for the level 1 ecoregion *r* containing route *s* is therefore a vector with 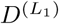 elements, given by:

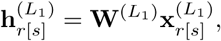

which essentially corresponds to extracting column *r* from the embedding matrix 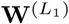.

Embeddings for route *s* are concatenated with the numeric route-level features **x**_*s*_ (a vector containing climate principal components, road density, distance from coast, latitude, and longitude) to obtain a vector for route *s* that is the zeroth hidden layer of the neural network 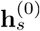:

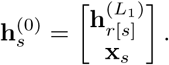

This vector 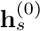 is mapped to next hidden layer with *D*^(1)^ = 4 hidden units via a fully-connected layer, followed by leaky rectified linear unit activation functions (Xu *et al*. 2015). The leaky rectified linear unit activation function is *g*(*x*) = max(0, *x*) − 0.01 × min(0, *x*). Thus, the next hidden layer is obtained in a forward pass via:

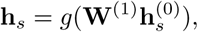

where **W**^(1)^ is a *D*^(1)^ × *D*^(0)^ parameter matrix. For the single species models, *D*^(0)^ = 21, because there is one embedding of size 8, and an additional 13 elements in the route-level covariate vector **x**_*s*_ (eight principal component axes, elevation, road density, distance from coast, latitude, and longitude).

The hidden layer is then mapped to initial occupancy, persistence, colonization, and detection probabilities, via fully connected layers:

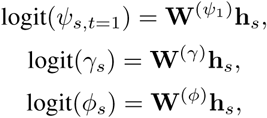

**Figure S3.1:**
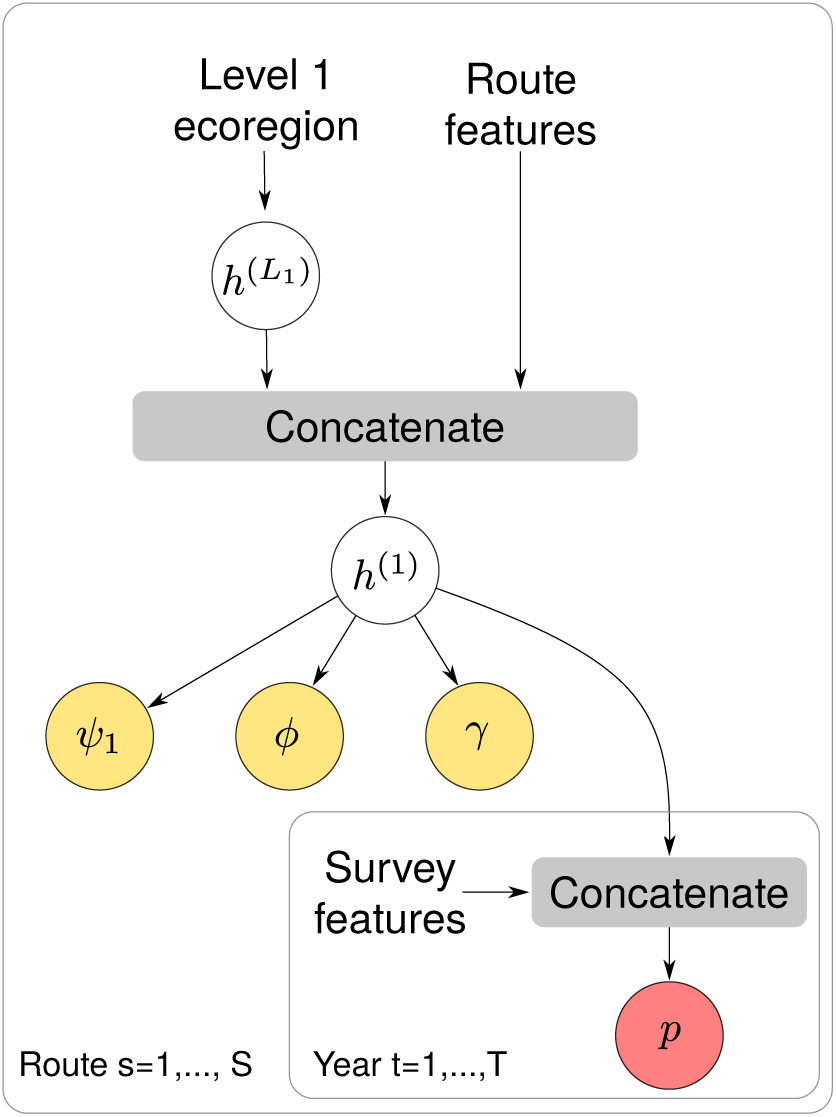
Extended computational diagram for the single species neural dynamic occupancy model. Outer grey boxes indicate the different levels of the model (route and year) that index quantities inside the boxes. Yellow nodes indicate occupancy parameters, and red nodes indicate detection parameters. Hidden layers are represented by *h*, with layer-specific superscripts. Outputs include initial occupancy (*ψ*_1_), persistence (*ϕ*), colonization (*γ*), and detection probabilities (*p*).

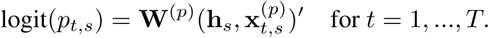

Here 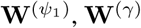, and **W**^(*ϕ*)^ are 4 × 1 parameter matrices, and **W**^(*p*)^ is a (4 + *n*_*p*_) × 1 parameter matrix that maps the concatenation of detection-related hidden unit vector 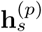 with *n*_*p*_ survey specific features contained in the vector 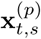 to the detection probability *p*_*t,s*_.

### Multi-species neural hierarchical model structure

#### Process model

The process model for the multi-species model extends the single-species models to the multiple species case. Presence/absence states *z*_*t,s,j*_ for all *s, t*, and *j* are equal to 0 if species *j* is absent from route *s* in time *t*, and *z*_*t,s,j*_ = 1 if the species is present. Occupancy dynamics are modeled as a function of initial occupancy, persistence, and colonization probabilities (Royle & Kéry 2007):

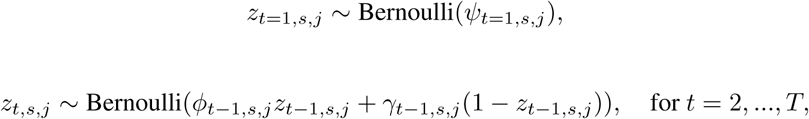

where *ψ*_*t,s,j*_ is the probability of occurrence of species *j* at route *s* in year *t, ϕ*_*t*−1,*s,j*_ is the probability of persistence conditional on presence in the previous year, and *γ*_*t*−1,*s*,,*j*_ is the probability of colonization conditional on absence in the previous year.

#### Observation model

The observation model similarly is a multi-species extension of the single species observation models. The observations *y*_*t,s,j*_ are integer-valued counts in the set 0, 1, 2, …, 50 that indicate the number of stops for which species *j* was detected in year *t* on route *s*. Detection/non-detection data are modeled as binomial random variables:

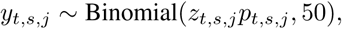

for all *s, t*, and *j*.

### Parameter model

#### Route representations

Heterogeneity in occupancy dynamics was introduced by allowing initial colonization, persistence, colonization, and detection probabilities to relate to latent spatiotemporal feature vectors (Fig. S3.2). These latent spatiotemporal feature vectors represent a nonlinear combination of route-level inputs. For the multi-species model, all feature vectors are 64 dimensional. Hidden layers are 32 dimensional unless stated otherwise.

Categorical inputs to the network included EPA level one ecoregion, mapped to categorical entity embeddings (Guo & Berkhahn 2016). As with the single-species models:

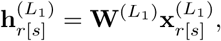

where 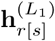 is the embedding for a level one ecoregion *r* containing route *s*, 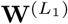 is a parameter matrix, and 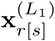 is a one-hot encoding for the level one ecoregion *r* containing route *s*. Concatenating the ecoregion embedding with route-level features **x**_*s*_ for route *s* provides the zeroth hidden layer of the network.

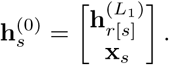

This zeroth hidden layer 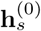 that combines an ecoregion embedding and route-level features is then passed to a sequence of hidden layers:

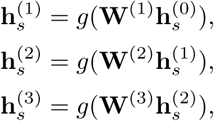

where *g* is the leaky ReLU activation function, so that 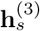 is a vector valued nonlinear combination of route-level features.

This hidden layer is then mapped to parameter-specific hidden layers. The initial occupancy probability hidden layer uses a fully connected layer:

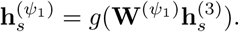

Colonization, persistence, and detection probability hidden layers were allowed to vary in space and time and modeled using a recurrent neural network. Temporal variation was modeled by treating 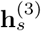 as a route-level encoding, that is decoded by a two-layer gated recurrent unit – a particular type of sequence model often used in time series analysis and text modeling, similar to a long short term memory model (Chung *et al*. 2014). The gated recurrent unit takes the vector 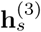 as an input, and outputs a multivariate sequence of shape *T* × (32 × 3), which is reshaped into a *T* × 32 × 3 dimensional array that has dimensions for years, hidden features, and model components (persistence, colonization, and detection).

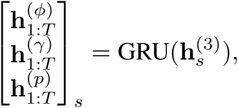

where the subscript 1: *T* indicates that values are generated for timesteps *t* = 1, …, *T*. These latent spatiotemporal route features are then combined with species-specific parameters generated via deep multi-species embedding (Chen *et al*. 2016).

#### Hierarchical deep multi-species embedding

Species-specific parameters were modeled as outputs of a neural network that ingests species-level traits (in this case, species identity, genus, family, and order). Taxonomic embeddings enable information to be shared among species within genera, families, and orders. A vector valued embedding **v**_*j*_ for species *j* is generated by concatenating embeddings at each taxonomic level:

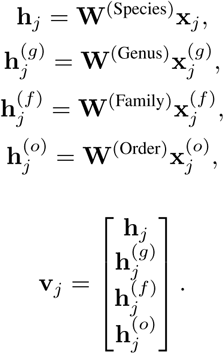

Here **h**_*j*_ is the vector valued embedding for species *j*, **W**^(Species)^ is a parameter matrix, **x**_*j*_ is a one hot encoded vector, 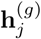 is the vector valued embedding for the genus containing species *j*, etc. The vector **v**_*j*_ is mapped to parameter-specific hidden units via fully connected layers:

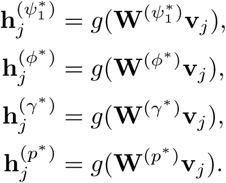

These hidden layers are then mapped to species-specific parameters via fully connected layers with identity activation functions:

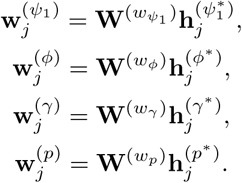

By modeling parameters as outputs of a neural network with shared features, relationships among occupancy and detection parameters can be learned. By including features (embeddings) at multiple taxonomic levels, information among taxonomically similar species can be shared.

#### Combining route features and species-specific parameters

**Figure S3.2:**
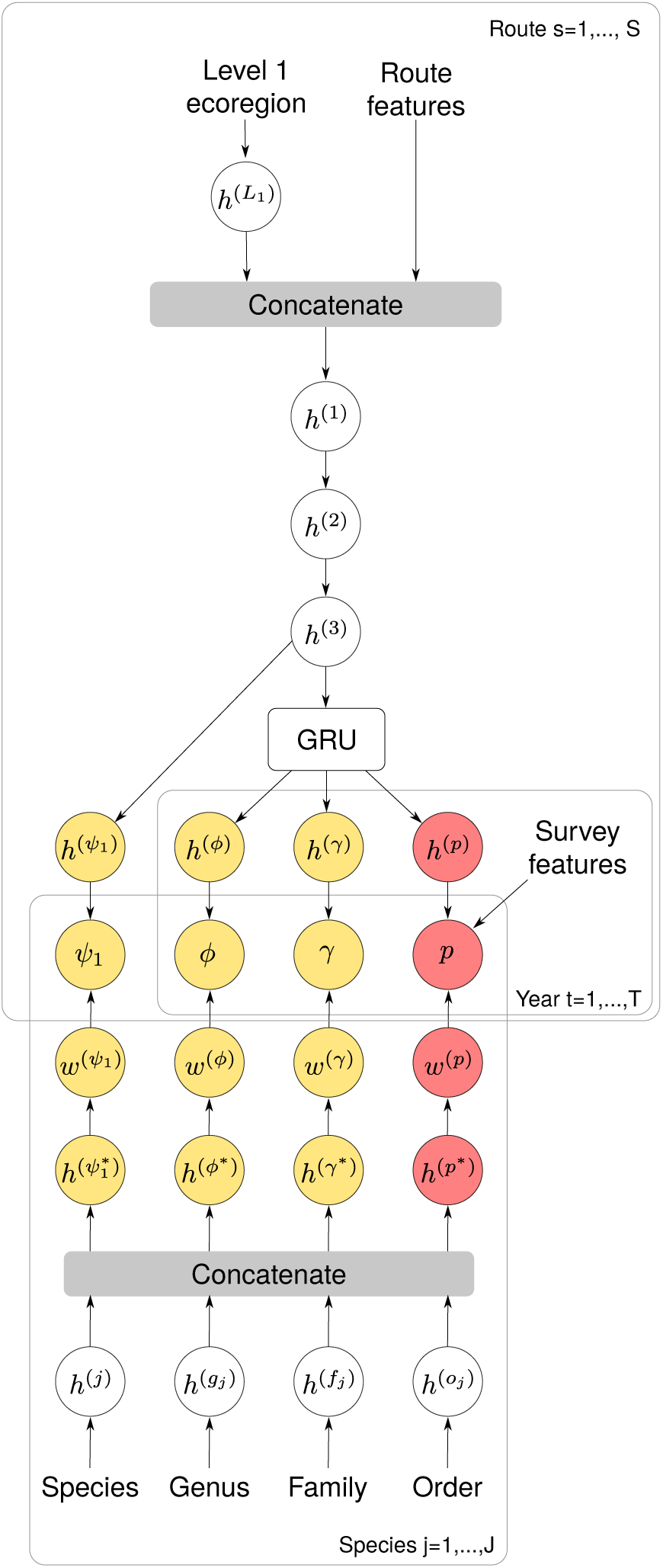
Extended computational diagram for the multi-species neural hierarchical dynamic occupancy model. Outer grey boxes indicate the different levels of the model that index quantities inside the boxes. Yellow nodes indicate occupancy parameters, and red nodes indicate detection parameters. Outputs include initial occupancy (*ψ*_1_), persistence (*ϕ*), colonization (*γ*), and detection probabilities (*p*). The box labeled GRU is a gated recurrent unit that decodes temporal sequences of hidden layers from encoded route vectors.

A dot product combines latent spatiotemporal route features with species-specific parameters:

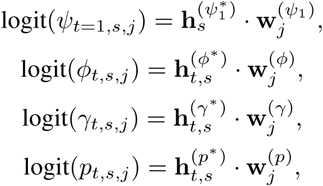

where **x** · **y** is the dot product of **x** and **y**: **x** · **y** = Σ_*i*_ *x*_*i*_*y*_*i*_.

#### Implementation

Maximum *a posteriori* estimates of baseline model parameters were obtained with Stan (Carpenter *et al*. 2017; Stan Development Team 2018). Penalized maximum likelihood estimates of the neural hierarchical model were obtained with PyTorch (Paszke *et al*. 2017), with L2 regularization penalties on the network parameters, which is equivalent to maximum a posteriori estimation for the neural network with Gaussian priors for the parameters (Blundell *et al*. 2015). The discrete latent occupancy states were marginalized using the forward algorithm, so that the optimization objective used the observed data likelihood (rather than the complete data likelihood). This optimization proceeded in one step, in contrast to some previous approaches that combine deep neural networks with spatiotemporal models using two-stage least squares parameter estimation (McDermott & Wikle 2019). To avoid underflow, forward probabilities were computed on the log scale for the baseline model in Stan, and using forward probability scaling in PyTorch for the neural hierarchical models (Rabiner 1989). All code required to reproduce the analysis is available on GitHub at https://www.github.com/mbjoseph/neuralecology.

